# *De Novo* Design of Miniprotein Inhibitors of Bacterial Adhesins

**DOI:** 10.1101/2025.08.18.670751

**Authors:** Adam M. Chazin-Gray, Tuscan R. Thompson, Edward D. B. Lopatto, Pearl Magala, Patrick W. Erickson, Andrew C. Hunt, Anna Manchenko, Pavel Aprikian, Veronika Tchesnokova, Irina Basova, Denise A. Sanick, Kevin O. Tamadonfar, Morgan R. Timm, Jerome S. Pinkner, Karen W. Dodson, Alex Kang, Emily Joyce, Asim K. Bera, Aaron J. Schmitz, Ali H. Ellebedy, Kelli L. Hvorecny, Gianluca Interlandi, Mark J. Cartwright, Andyna Vernet, Sarai Bardales, Desmond White, Rachel E. Klevit, Evgeni V. Sokurenko, Scott J. Hultgren, David Baker

## Abstract

The rise of multidrug-resistant bacterial infections necessitates the discovery of novel antimicrobial strategies. Here, we show that protein design provides a generalizable means of generating new antimicrobials by neutralizing the function of bacterial adhesins, which are virulence factors critical in host-pathogen interactions. We *de novo* designed high-affinity miniprotein binders to FimH and Abp chaperone usher pili adhesins from uropathogenic *Escherichia coli* and *Acinetobacter baumannii*, respectively, which are implicated in mediating both uncomplicated and catheter-associated urinary tract infections (UTI) responsible for significant morbidity worldwide. The designed antagonists have high specificity and stability, disrupt bacterial recognition of host receptors, block biofilm formation, and are effective in treating and preventing murine models of uncomplicated and catheter-associated UTIs *in vivo*.

## Introduction

Multidrug-resistant (MDR) bacterial pathogens constitute a significant and growing threat to public health [1]. To counter the alarming rise of these increasingly untreatable and deadly bacterial infections, accelerating the development of novel antimicrobial therapies is essential. Recent advances in protein design have opened the door to designing small (< 120 amino acids) *de novo* protein binders (minibinders) that precisely target virtually any epitope [2, 3], enabling therapeutics against viruses [4], toxins [5], and other disease targets [6]. However, a major challenge in applying these tools to combat bacteria is the accessibility of minbinders to intracellular and membrane-embedded target antigens due to their shielding by surface molecules such as lipopolysaccharides. Previous attempts have identified high-affinity protein binders to their integral outer membrane protein target antigens *in vitro*, only to find that the binders cannot access their targets in a cellular context [7, 8].

We reasoned that bacterial adhesins would constitute ideal targets for designed minibinder therapeutics targeting bacteria because they lie well beyond bacterial surface molecules and play a key role in the virulence of MDR bacteria by mediating their attachment to and invasion of host epithelial cells as well as biofilm formation (Fig. 1a). Among the diverse family of bacterial adhesins, chaperone-usher pathway (CUP) pili adhesins lie at the tips of extracellular fibers and recognize receptors in a lectin-substrate interaction that mediates host tropisms. CUP pili are ubiquitous among gram-negative bacteria and are critical for the establishment and persistence of numerous bacterial infections, including both uncomplicated urinary tract infections (UTIs) and catheter-associated UTIs (CAUTIs). UTIs are one of the most common bacterial infections among women and their treatment accounts for approximately 15% of antibiotic usage in the United States [9, 10]. Over 50% of UTIs, especially those acquired in hospitals, are caused by MDR bacteria, limiting the effectiveness of a dwindling number of traditional antibiotic treatments [11, 12, 13]. Previous anti-adhesin therapeutic strategies have relied on small molecule ligand mimetic inhibitors [12] or monoclonal antibodies (mAbs) [13, 15]. However, these strategies require identification of the adhesin substrate, which is often unknown, or labor-intensive screening for inhibitory mAbs. Designed minibinders could overcome this therapeutic bottleneck by enabling the rapid development of antibiotic-sparing therapies, relying solely on the adhesin structure and identification of its putative binding pocket. To investigate this, we sought to design minibinders against three CUP adhesins: FimH from *Escherichia coli*, which accounts for 70-90% of UTIs, and Abp1D and Abp2D from *Acinetobacter baumannii*, which is prevalent in hospital-acquired CAUTIs and is often multidrug resistant.

**Fig. 1.**
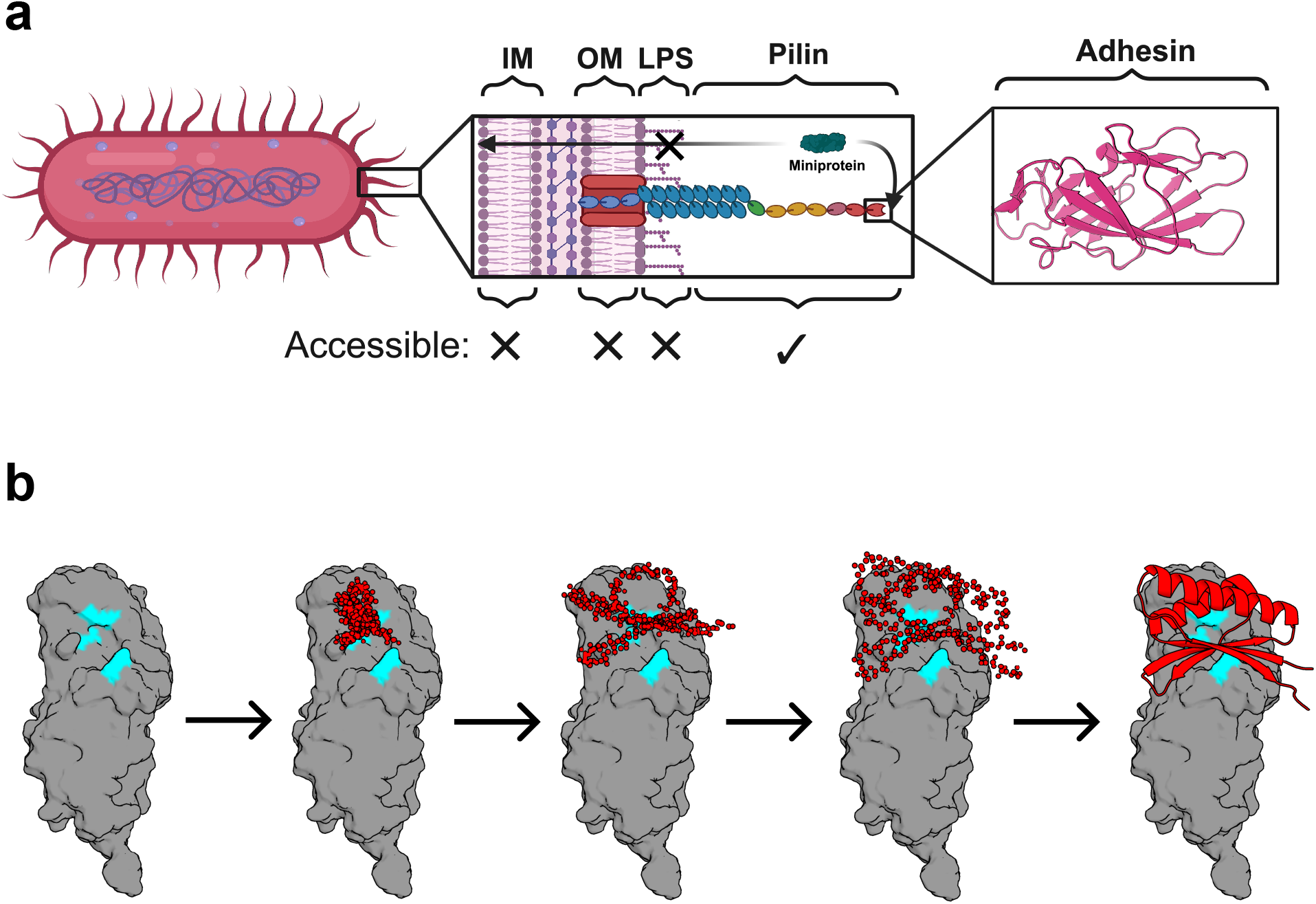
*De novo* design of adhesin inhibitors. **(A)** Schematic illustrating the structures on the gram-negative bacterial cell surface that are potentially accessible to protein-based inhibitors. Virulence factors that make up the bacterial pili, especially tip adhesins, are attractive targets due to their essential role in pathogenesis and the fact that they lie well off the membrane and beyond the protective lipopolysaccharide shell. Created in BioRender. **(B)** Example denoising trajectory for an Abp2D binder. “Hotspots” corresponding to the adhesin substrate binding pocket residues are highlighted in cyan.

## Results

### *De novo* design of minibinders targeting FimH

Uropathogenic *E. coli* (UPEC) expresses type 1 pili, a prototypical CUP pilus, which are tipped with the mannose binding adhesin FimH [14, 13, 15]. FimH is highly invariant with greater than 90% amino acid conservation among *E. coli* isolates (sequence provided in Supplementary Data File 1). Further, the mannose binding pocket of FimH is invariant amongst all *Enterobacteriaceae* [15, 16]. UPEC FimH is essential for bacterial attachment to the bladder epithelium via mannosylated glycoproteins [17]. After attachment, UPEC forms intracellular bacterial communities that are recalcitrant to antibiotics and, if left untreated, can lead to recurrent UTIs and kidney infection [17]. FimH consists of an N-terminal mannose-binding lectin domain and a C-terminal pilin domain that attaches to the rest of the pilus [14, 17]. The lectin domain is an allosteric protein that samples a conformational equilibrium between two well-characterized conformational states: a low-affinity state (LAS, also known as the “Tense” state) and a high-affinity state (HAS, also known as the “Relaxed” state) [18]. In the LAS, the body of the FimH lectin domain interacts with the pilin domain, fixing it in place, and allosterically disrupts the mannose binding pocket, resulting in a low affinity for mannose [14, 17]. In the HAS, the lectin domain has little to no interaction with the pilin domain, allowing it to rotate freely relative to the pilin domain, and the mannose binding pocket loops are stably positioned to bind to mannose [17].

We used RFdiffusion [3] to generate novel minibinder backbones against both states of FimH. We targeted designs to the binding pocket of published crystal structures (FimH in LAS: 3JWN; FimH in HAS: 1UWF) by specifying “hotspot” residues in the binding pocket as inputs to the model (Fig. 1b). We subsequently assigned sequences to these backbones using ProteinMPNN [19] and scored designs with AlphaFold2 (AF2) Initial Guess [20]. The best sequences ranked by AF2 folding and interaction confidence metrics (predicted local distance difference test (pLDDT) and predicted aligned error (pAE) scores) were then iteratively refined by partially diffusing [21] the AF2 predicted structures with RFdiffusion, reassigning sequences with ProteinMPNN, and filtering with AF2. We obtained an oligonucleotide array encoding the top 10,000 designs by AF2 and Rosetta interface metrics.

### Experimental characterization of FimH minibinders

FimH minibinders were screened via cDNA display, rather than a cell-based screening approach like yeast surface display, because FimH binds to mannose on the cell surface. We panned against the wild-type FimH lectin domain, which adopts the HAS, and a mutant lectin domain stabilized in the LAS [22]. The top 96 designs enriched more than four-fold over the naive library and consisted of 78 binders identified through the HAS sorts and 39 binders identified through the LAS sorts (raw sequencing data provided in the SRA database under BioSample accession numbers: SAMN62230579-SAMN62230585 [http://www.ncbi.nlm.nih.gov/bioproject/1508338]). Twenty-one designs were enriched against both states. Eighty-one of the enriched sequences were designed against the LAS.

We expressed these 96 FimH minibinders in *E. coli* and found that 79 exhibited monodisperse size exclusion chromatography (SEC) traces and expressed sufficiently for downstream characterization (see Materials and Methods for details). In an initial surface plasmon resonance (SPR) binding screen, nine of these designs exhibited sub-micromolar binding against at least one of the FimH states (table S1). All nine designs were originally designed against the LAS. To directly assess the ability of FimH minibinders to inhibit and disrupt bacterial adhesion, we performed a series of red blood cell (RBC) agglutination (or hemagglutination) assays using *E. coli* strain KB23 expressing type 1 fimbriae. *E. coli* agglutinates RBCs by binding to mannosylated glycoproteins on their surface via FimH. Of the 96 minibinders we tested, eight significantly inhibited hemagglutination (fig. S1A), seven of which also had sub micromolar affinities in the SPR screen. Minibinder F7 had the lowest minimum inhibitory concentration (69 nM) in the RBC agglutination inhibition assay and was therefore selected for further characterization (fig. S1B, table S2). F7 was designed against the LAS, expressed well in *E. coli*, and purified as a monodisperse species (Fig. 2a, 2b). Though it was designed against the LAS, F7 was identified by panning against the HAS. Further characterization by SPR confirmed that F7 binds to both conformations, but with a several-fold higher affinity to the LAS (K_d_=119 nM) than HAS (K_d_=713 nM) (Fig. 2c). F7 exhibited the expected circular dichroism (CD) spectra and proved to be highly thermostable, with a melting point above 75°C (Fig. 2d, 2e).

**Fig. 2.**
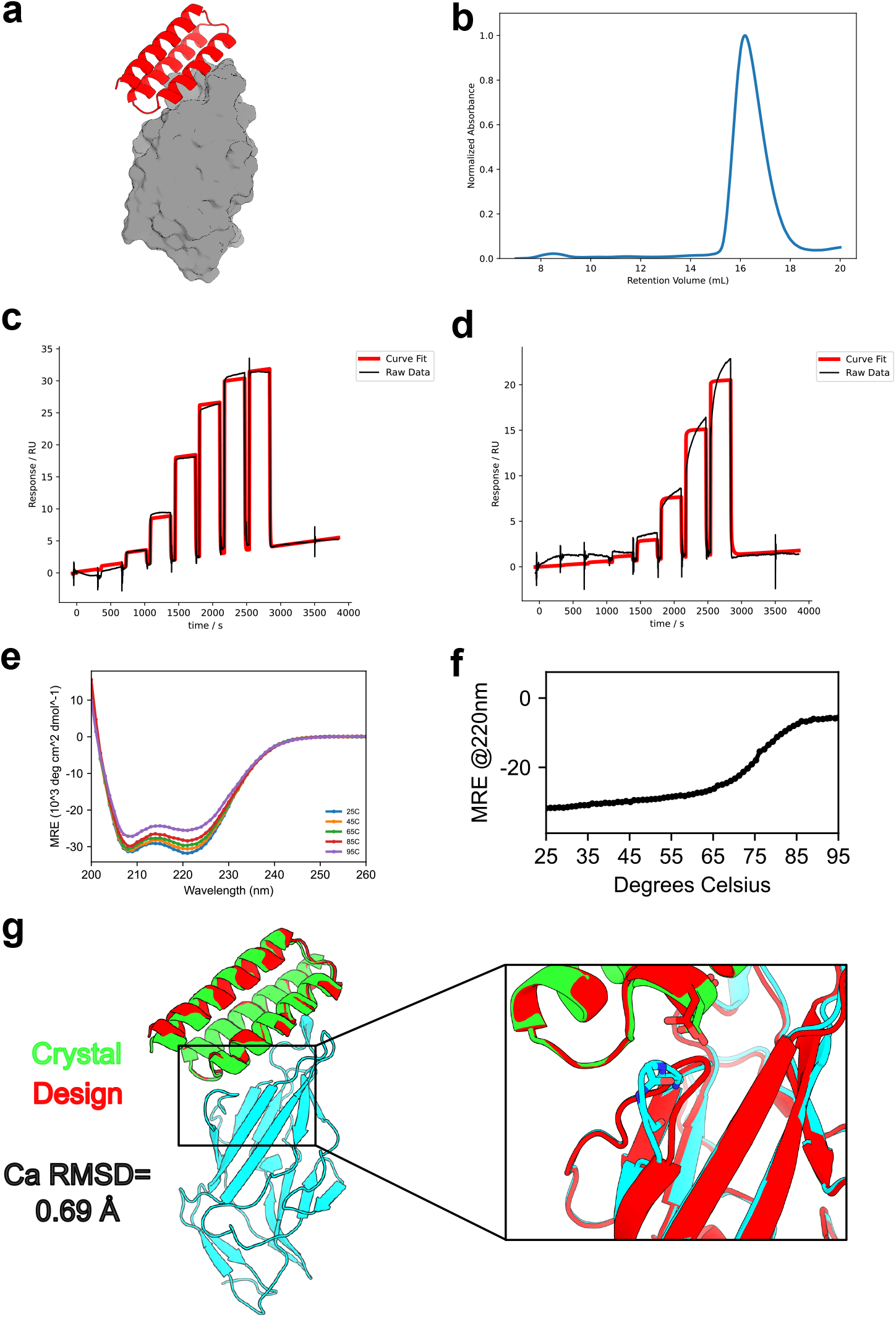
*In vitro* characterization of FimH minibinder F7. **(A)** Design model for FimH minibinder F7 (red) in complex with FimH LAS (gray). **(B)** SEC trace of F7: sample was injected onto a Superdex 75 10/300 GL column. **(C)** The binding affinity of F7 was determined with SPR. The SPR data indicate that F7 binds more tightly to the LAS (left; K_d_ =119 nM) than HAS (right; K_d_ =713 nM). Raw data (black) is overlaid with fitted data (red). **(D)** Raw CD data for F7. Full wavelength scans from 250 to 200 nm were performed at the indicated temperatures (25°C: blue, 45°C: orange, 65°C: green, 85°C: red, 95°C: purple). **(E)** A melting curve for F7 indicates that it is thermostable up to 75°C. CD signal at 220 nm (helicity) was measured every 1°C. **(F)** Crystal structure of F7 (green) with FimH L34K (cyan) overlaid on the design model (red); inset: difference in FimH clamp loop between design and crystal structure showing an unexpected displacement of the clamp loop. The experimental structure exhibits a Ca RMSD of 0.69 Å to the design model. Source data are provided as a Source Data file.

Co-crystallization of F7 with a mutant FimH lectin domain stabilized in the LAS (L34K; sequence provided in Supplementary Data File 1)[23] confirmed the designed binding interface with a C*a* RMSD of 0.69 Å to the design model (PDB Code: 9Q1V; Fig. 2f, fig. S2, table S3). The experimentally determined structure of FimH bound to F7 was much closer to the canonical LAS conformation than the HAS conformation (C*a* RMSD of 0.45 Å vs 2.26 Å). As expected, the minibinder occupies the mannose-binding pocket of FimH (fig. S3). [24] Unexpectedly, it also wedged between one of the mannose-binding loops and the clamp loop, further displacing it from the binding pocket compared to the design model. The clamp-loop displacement induced by F7 closely resembles that of the recently determined structure of FimH bound to anti-FimH antibody mAb926, which was raised against the LAS and known to function through a parasteric mechanism (fig. S4). [13]

Given that F7 preferentially binds the LAS-stabilized mutant FimH and that the crystal structure of FimH in complex with the minibinder resembles the LAS, we suspected that F7 binding may induce a shift from the HAS to the LAS in wild-type FimH lectin domain. We compared F7’s interaction with the L34K variant of the lectin domain (LAS) and wild-type lectin domain (HAS) by (^15^N, ^1^H)-HSQC NMR spectroscopy. The substantial structural differences between the LAS and HAS conformations of the isolated lectin domain are reflected in their distinct NMR spectra, which display very little peak overlap (fig. S5A, S5B). Addition of F7 to the LAS variant caused chemical shift perturbations localized at the binding interface, consistent with a binding event and/or local change in conformation, but not a global conformational change (fig. S5C, S6B). In contrast, the spectrum of the WT HAS domain showed widespread chemical shift changes in the presence of F7, becoming similar to that of the LAS variant (fig. S5C, S6A). Notably, spectra of the F7-bound HAS and LAS variants now show extensive peak overlap, indicating a shared conformation (fig. S5C). Together, these data indicate that F7 binding to the isolated WT lectin domain induces a conformational change from the HAS to the LAS, shifting the conformational equilibrium in the opposite direction as binding the native ligand and existing mannoside therapeutics. [18].

We quantitatively assessed the ability of minibinder F7 to block bacterial adhesion to surfaces coated with bovine RNAseB—a model glycoprotein rich in Man_5_ high-mannose type oligosaccharides that avidly binds FimH even in the LAS. We preincubated *E. coli* KB23 with either F7, alpha-methyl-mannoside, mAb926, or a minibinder that showed no inhibition in hemagglutination assays (A4) and then measured its binding to RNaseB. F7 displayed an IC50 of 1.9 µM (95% CI: 1.4-2.9 µM), a more than two-thousand-fold improvement over mannose (4.7 mM) but had a higher IC50 than mAb926 (40 nM; Fig. 3a). We then performed a detachment ELISA by first adhering bacteria to RNAseB-coated plates and subsequently introducing either F7, noninhibitory minibinder A4, or mAb926, which is known to effectively dissolve preformed biofilms. [25] F7, but not A4, successfully detached UPEC with an IC50 of 1.7 µM (95% CI: 1.3-2.4 µM), but had a greater IC50 than mAb926 (9 nM) by two orders of magnitude (Fig. 3b). We also sought to characterize F7’s inhibitory ability in a series of more biologically relevant RBC assays. Unlike A4, minibinder F7 inhibited RBC agglutination by *E. coli* with a minimum inhibitory concentration of 78 nM (Fig. 3c), which was lower than alpha-methyl-mannoside by more than a thousand-fold (140 µM; fig. S7) but higher than that of mAb926 by ~100-fold (0.1 nM; Fig. 3c). We tested the ability of F7, mAb926, and A4 to reverse bacterial hemagglutination by adding it to pre-agglutinated mixtures of RBC and *E. coli*. Unlike minibinder A4, F7 and mAb926 successfully redispersed the RBCs within 15 minutes of introduction at 1 µg/ml (Fig. 3d). We confirmed the physiological relevance of F7’s inhibitory ability by comparing the adherence of UPEC to T24 bladder cells in the presence and absence of minibinder using microscopy. F7-treated cells had significantly fewer bacteria (mean = 2.85, SEM+/−0.57) per T24 cell present than either buffer-treated cells (mean = 17.26, SEM+/−2.24, p=0.0012, ANOVA) or cells treated with A4 (mean = 29.79, SEM+/−4.86, p<0.0001, ANOVA) but had no significant difference from mAb926 treated cells (mean = 2.0, SEM+/−0.42, n.s., ANOVA, Fig. 3e, fig. S8).

**Fig. 3.**
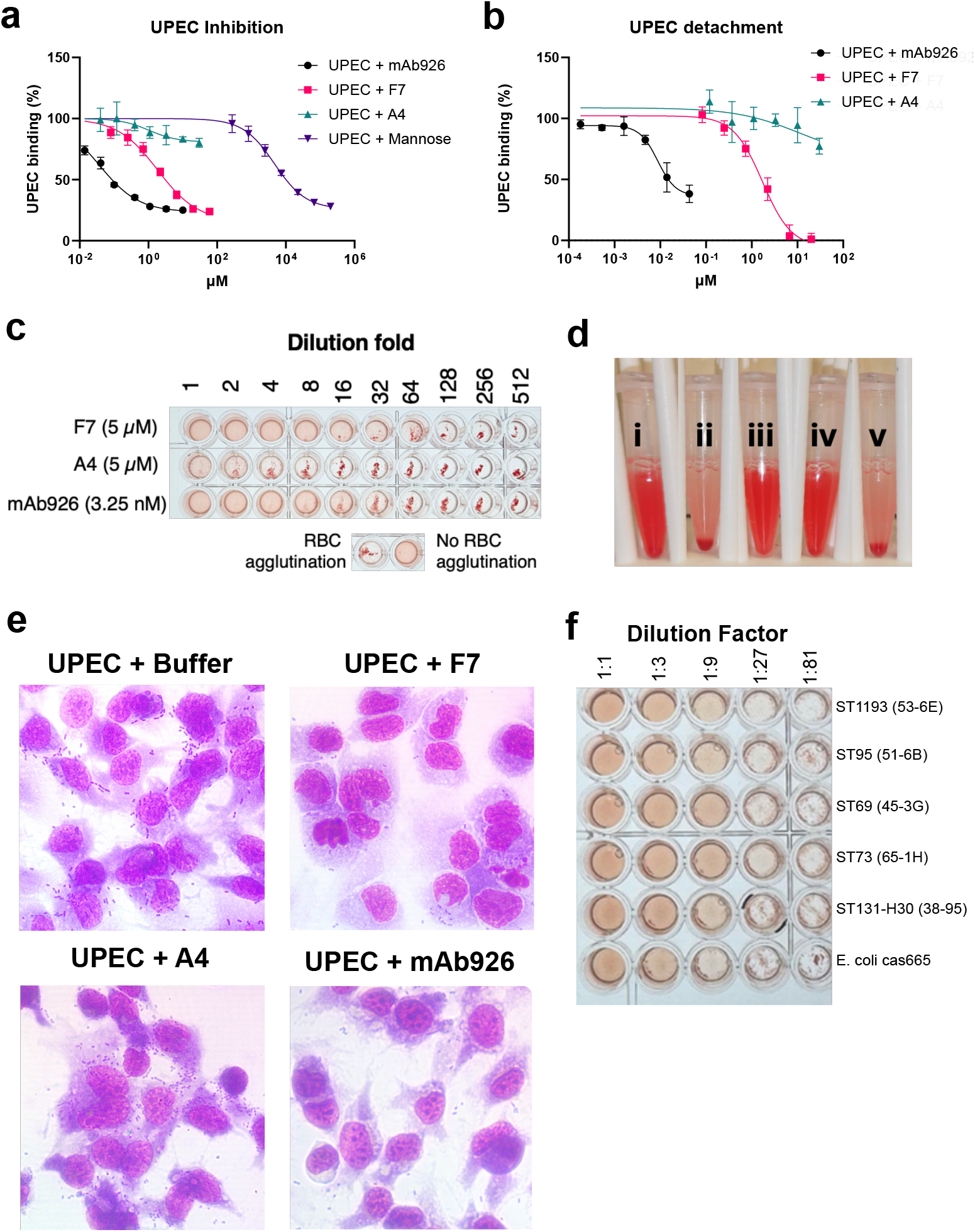
*In cellulo* characterization of FimH minibinder F7. **(A)** Inhibition ELISA results for minibinder F7 (pink; IC50=1.9 μM; 95% CI: 1.4-2.9 μM), noninhibitory minibinder A4 (teal), mAb926 (black; IC50=40 nM), and mannose (purple; IC50=4.7 mM). Each experiment includes three biological replicates. Data are presented as mean values +/− standard deviation (SD). **(B)** Detachment ELISA results for F7 (pink; IC50=1.7 μM; 95% CI: 1.3-2.4 M), A4 (teal), and mAb926 (black; IC50=9 nM). Each experiment includes three biological replicates. Data are presented as mean values +/− SD. **(C)** Red blood cell agglutination inhibition assay with a serial dilution of F7, A4, and mAb926. **(D)** Red blood cell agglutination dispersal assay showing red blood cells incubated for one hour without bacteria (i), with bacteria (ii), with bacteria & mAb926 (iii), with bacteria & minibinder F7 (iv), and with bacteria & minibinder A4 (v). **(E)** Representative microscopy images of T24 bladder cells incubated with UPEC and treated with miniprotein F7, mAb926, miniprotein A4 or buffer. The addition of F7 and mAb926 decreases the number of bacteria present. Images are shown at 20x magnification. These images are representative of 3 independent experiments. **(F)** F7 inhibition of RBC agglutination by clinical *E. coli* strains from the clonal groups (STs) of major clinical importance and *E. coli* cas665 expressing *K. pneumoniae* FimH. Source data are provided as a Source Data file.

Given the sequence diversity of FimH across disease-causing strains, we tested whether F7 is inhibitory not only to the FimH variant it was designed against but also to different FimH variants expressed by the most common highly-virulent and/or multi-drug resistant clonal groups of UPEC. F7 successfully inhibited hemagglutination by strains from the clonal groups ST73, ST95, ST69, and ST131-H30 (Fig. 3f, sequences provided in Supplementary Data File 1). F7 also inhibited agglutination by *E. coli* strain cas665 expressing *Klebsiella pneumoniae* FimH, which has approximately 85% sequence identity to *E. coli* FimH, suggesting that F7 may be cross-reactive with FimH variants from other bacterial pathogens (Fig. 3f; sequence provided in Supplementary Data File 1).

### *De novo* design of minibinders targeting Abp adhesins

CAUTIs caused by MDR bacteria pose a significant economic toll on the healthcare system in the United States.[26] CAUTIs represent approximately 40% of hospital-acquired infections and have an increased risk of morbidity and mortality compared to uncomplicated UTIs.[26, 27] CAUTIs result in a serious deterioration of quality of life through pain, discomfort, disruption of daily activities, and increased healthcare costs, exacerbated by the rapid spread of drug resistance in uropathogens. [26, 27, 28, 29, 30]

One particularly concerning CAUTI pathogen is *A. baumannii*, which is classified by the CDC as a pathogen of urgent concern due to its high rates of multidrug resistance and increases in relative prevalence in CAUTIs compared to uncomplicated UTIs.[31] The *A. baumannii* strain “ACICU” possesses two highly similar CUP pili, Abp1 and Abp2, that play critical roles in mediating CAUTI. The Abp1 and Abp2 pili are tipped with the Abp1D and Abp2D adhesins that allow these bacteria to bind to host-deposited fibrinogen on implanted catheters, which is a critical step for *A. baumannii* to establish a CAUTI.[32] Approximately 63% of all *A. baumannii* strains contain either an Abp1 or Abp2 pilus in their genome.[32] Among genomes of clinical *A. baumannii* isolates analyzed, over 90% contained *abp2D*, with the Abp2D allele present in strain ACICU representing the predominant protein variant, occurring in over 54% of isolates encoding the *abp2* operon.[33]

While the receptor binding domains of the ACICU Abp1D and Abp2D adhesins share only 70% sequence identity, they share a higher degree of structural homology as seen in previously solved crystal structures (Abp1D: 8DF0; Abp2D: 8DEZ; sequences provided in Supplementary Data File 1) [32]. Previous structure-function studies identified the putative fibrinogen-binding pockets of these adhesins using mutagenesis; however, the exact structural binding interactions between fibrinogen and the adhesins remain unknown.[32] Similar to the dynamic conformational ensembles displayed by FimH, the binding affinity of the *A. baumannii* Abp1D and Abp2D receptor binding domains (RBD) are controlled via dynamics of the flexible anterior binding loop[32], illustrated by comparing the loop’s open low-affinity conformation in the Abp2D crystal structure to an AlphaFold2 model for this protein displaying the loop closed in its high-affinity conformation (fig. S9). In addition, while vaccination with Abp2D protected mice from *A. baumannii* CAUTI, these efforts to immunize mice with Abp2D failed to identify sufficiently inhibitory mAbs to effectively disrupt binding of ACICU to fibrinogen (IC50s > 100 µM) in a bacterial ELISA assay (fig. S10).[33] Due to the promise of anti-adhesin therapeutics and the previous failed attempts at antibody identification, we aimed to apply protein design tools to develop more effective Abp inhibitors.[34]

We used RFdiffusion [3] with “hotspot” conditioning to generate novel minibinder backbones specifically targeting the putative fibrinogen-binding sites of both the ACICU Abp1D and Abp2D adhesins (Fig. 1b). Taking into account the anterior loop conformational differences between the high-affinity and low-affinity states, we designed backbones against the crystal structures of Abp1D and Abp2D receptor binding domains, which display the anterior binding loop in the open low-affinity conformation, exposing the putative binding pocket. We assigned sequences to these backbones using ProteinMPNN, predicted their structures with AF2 Initial Guess, and then iteratively optimized high-quality designs as described above. The top ~5000 scoring designs against Abp1D (~1300) and Abp2D (~3700), with a maximum of 15 sequences per backbone, were obtained as a DNA oligo chip.

### Experimental characterization of Abp minibinders

Abp adhesins do not bind mannose and are therefore amenable to screening minibinders using yeast surface display [4, 35]. Designed Abp binder sequences were amplified from the DNA oligo chip and cloned into a yeast display expression vector using yeast cloning. The resulting yeast display library was enriched for binding via two rounds of fluorescence-activated cell sorting (FACS) with 1 µM biotinylated Abp1D or Abp2D lectin domains using PE-conjugated streptavidin as a detection reagent. A third round of FACS sorting involved staining the enriched populations with 10-fold titrations of these adhesins from 1000 nM to 1 nM, revealing strong binding signal for the Abp2D-enriched population down to 1 nM, and binding signal down to 10 nM for the Abp1D-enriched population (fig. S11, Supplementary Data File 2). Next-generation sequencing of sorted populations identified 31 highly enriched binder sequences across both populations, including some sequences that were enriched in both Abp1D and Abp2D sorts, suggesting that we identified cross-reactive binders (sequencing counts are provided in Supplementary Data File 3). Interestingly, all 31 of these enriched binders were designed using the Abp2D crystal structure, which might be explained by the presence of a citrate molecule in the Abp2D crystal structure that opens its putative binding pocket and exposes more hydrophobic residues than in the Abp1D structure.

Minibinder sequences that were enriched during yeast surface display with Abp adhesins were expressed in and purified from *E. coli*. Of the 31 enriched binders, twenty-four exhibited primarily monodisperse SEC traces and had sufficient yields for downstream characterization. An initial SPR binding screen of these 24 binders against both Abp1D and Abp2D identified 16 binders that exhibited sub-micromolar affinity to Abp2D, four of which also exhibited sub-micromolar affinity to Abp1D (table S4). A follow-up SPR experiment on the four cross-reactive binders revealed that Abp minibinder A7 (design model in Fig. 4a) exhibited high affinity to both Abp1D (K_d_=50.4 nM) and Abp2D (K_d_=3.5 nM; Fig. 4c, table S2). A7 expressed well (~50 mg/L of culture), behaved as a monodisperse species as assessed by SEC (Fig. 4b), exhibited the expected CD spectra (Fig. 4d), and was thermostable with a melting temperature greater than 85°C (Fig. 4e). Next, we confirmed the specificity of the designed adhesin binders using SPR. Minibinder A7 exhibited undetectable levels of binding to FimH LAS or HAS (fig. S12). Similarly, FimH minibinder F7 exhibited insignificant binding to Abp1D and Abp2D (fig. S12).

**Fig. 4.**
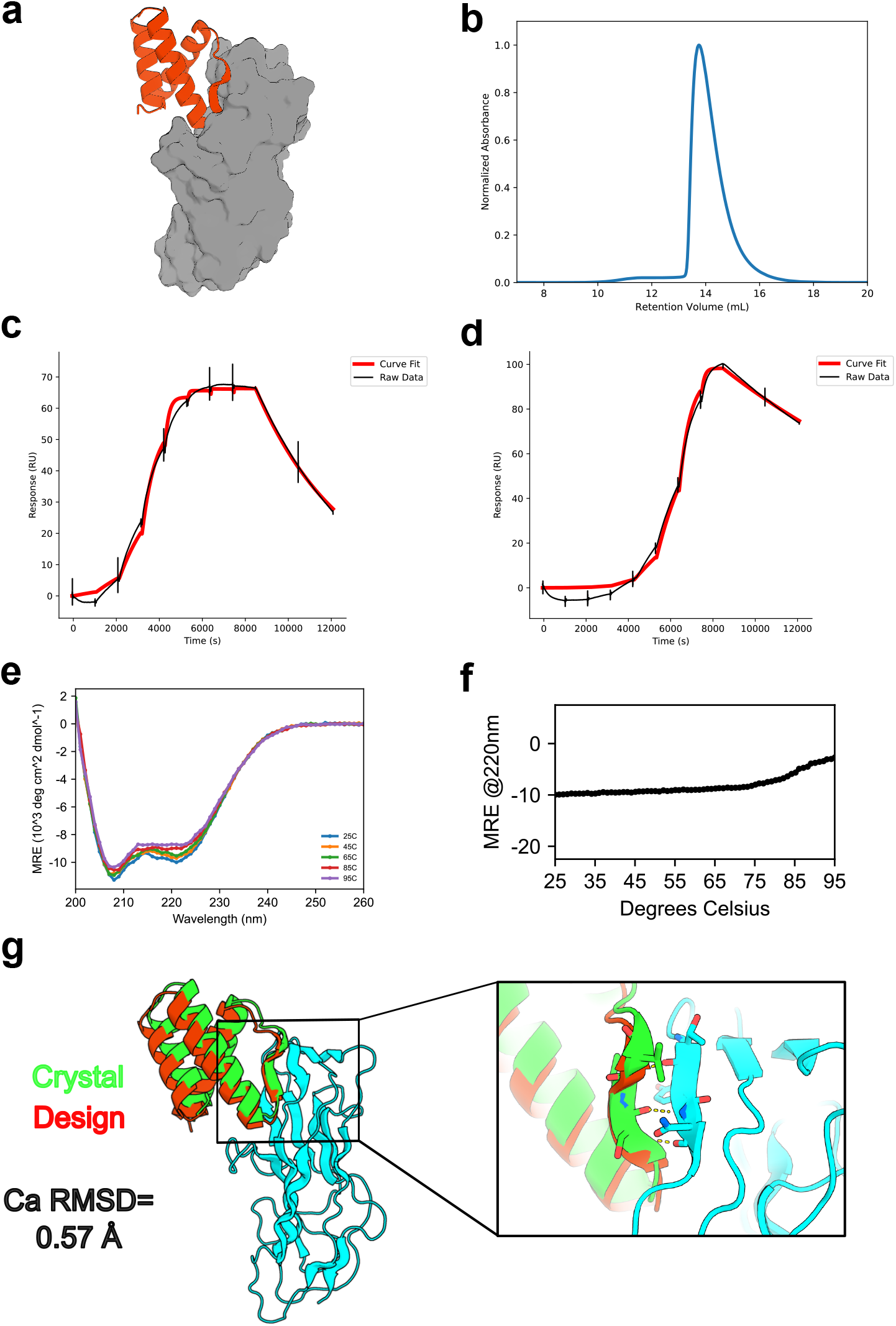
*In vitro* characterization of Abp minibinder A7. **(A)** Design model for Abp minibinder A7 (red) in complex with Abp2D (gray). **(B)** SEC trace of A7: sample was injected onto a Superdex 75 10/300 GL column. (**C**) The binding affinity of A7 was determined with SPR. The SPR data indicate that A7 binds with high affinity to Abp1D (left; K_d_ =50.4 nM) and Abp2D (right; K_d_ =3.5 nM). Raw data (black) is overlaid with fitted data (red). Raw CD data for A7. Full wavelength scans from 260 to 200 nm were performed at the indicated temperatures (25°C: blue, 45°C: orange, 65°C: green, 85°C: red, 95°C: purple). **(E)** A melting curve for A7 indicates that it is thermostable up to 80°C. CD signal at 220 nm (helicity) was measured every 1°. **(F)** Crystal structure of A7 (green) with Abp2D (cyan) overlaid on the design model (red); inset: beta-strand pairing interaction between A7 and an edge strand of Abp2D. The experimental structure exhibits a Ca RMSD of 0.88 Å to the design model. Source data are provided as a Source Data file.

Determining the structure of a co-crystal of A7 with Abp2D confirmed the designed interface, and exhibited a C*a* RMSD of 0.88 Å to the design model (PDB Code: 9Q1H; Fig. 4f, fig. S13, table S3). This structure confirms the designed binding mode that utilizes a helix-turn-strand motif to simultaneously stabilize the core of the minibinder, insert a loop deep into the putative fibrinogen binding pocket of Abp2D, and strand pair with an edge strand of this adhesin. As designed, A7 stabilizes the anterior binding loop of Abp2D in an open conformation, thus neutralizing the adhesin in the low-affinity conformation and matching the loop conformation in the Abp2D crystal structure.

To assess the ability of A7 to inhibit bacterial adhesion, we performed a bacterial ELISA assay with *A. baumannii* ACICU. When preincubated with ACICU prior to being added to the fibrinogen-coated ELISA plate, A7 but not a BSA control significantly inhibited ACICU binding with an IC50 of 2.6 nM (95% CI: 1.8-3.0 nM; Fig. 5a). In a “detachment” ELISA, A7 successfully displaced adherent ACICU and exhibited an IC50 of 40.0 nM (95% CI: 25.5-52.1 nM; Fig. 5b). Since bacterial adhesins are known to contribute to biofilm formation [32], we also tested whether designed inhibitors could both prevent ACICU from forming biofilms and disperse preformed biofilms. Crystal violet staining of *A. baumannii* biofilms grown on PVC plates demonstrated that 1 µM of A7 could prevent the formation of ACICU biofilms, unlike a BSA control (p < 0.0001, one-way ANOVA; Fig. 5c). In addition, 1 µM of A7 but not a BSA control could almost completely disperse preformed ACICU biofilms after 2 hours of treatment (p=0.0021, one-way ANOVA), mimicking the phenotype of an ACICU Δabp1 Δabp2 double knockout strain (Fig. 5c; p=0.0003, one-way ANOVA).

**Fig. 5.**
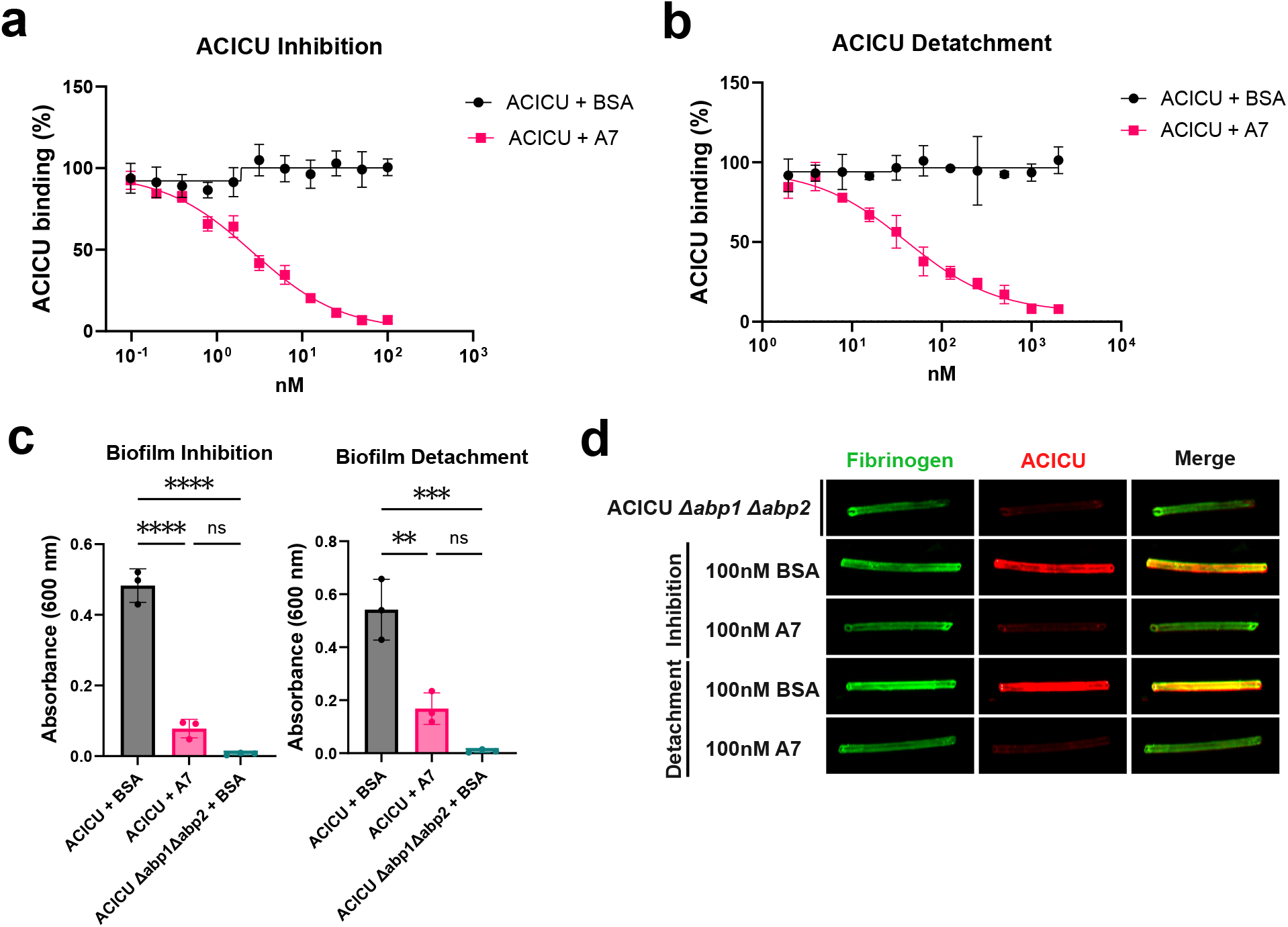
*In cellulo* characterization of Abp minibinder A7. **(A)** Inhibition ELISA results for minibinder A7 (pink; IC50=2.6 nM; 95% CI: 1.8-3.0 nM) and BSA (black) *A. baumannii* ACICU treatment groups. Each experiment includes three biological replicates. Data are presented as mean values +/− SD. **(B)** Detachment ELISA results for A7 (pink; IC50=40.0 nM; 95% CI: 25.5-52.1 nM) and BSA (black) *A. baumannii* ACICU treatment groups. Each experiment includes three biological replicates. Data are presented as mean values +/− SD. **(C)** A7 (pink) significantly inhibits the formation of biofilms (left, p<0.0001) and disperses existing biofilms (right; p=0.0021) relative to a BSA (grey) treatment control (n=3 independent biological replicates). Similarly, ACICU Δabp1 Δabp2 treated with BSA (teal) exhibits significantly lower levels of biofilm formation than the WT ACICU strain treated with BSA in both the inhibition (p<0.0001) and detatchment (p=0.0003) experiments. Data are presented as mean values +/− SD. ne-way ANOVA with Tukey’s multiple comparisons test. ****P ≤ 0.0001, ***P ≤ 0.001, **P ≤ 0.01, *P ≤ 0.05. **(D)** Fluorescence imaging of fibrinogen-coated silicone tubing reveals differential adherence of *A. baumannii* ACICU after pretreatment with or dispersion by A7 or a BSA control. The catheters in this panel were captured in a single imaging session and uniform processing was applied to the entire image to assess staining of Fibrinogen (green) and bacteria (red). Image representative of 4 independent replicates (except for 100 nM BSA detachment; n=3). Source data are provided as a Source Data file.

To explore whether minibinder A7 could disrupt *A. baumannii* ACICU biofilms in a more *in vivo*-like context, inhibition and detachment studies were performed with fibrinogen-coated silicone catheters.[32, 36] After coating silicone tubing with fibrinogen, ACICU was allowed to bind in the presence of A7 or BSA. We also tested the ability of A7 and BSA to disperse ACICU that had already bound to the silicone catheter. Immunostaining the tubing revealed that despite fibrinogen coating each piece of tubing, 100 nM of A7 but not the BSA control significantly prevented ACICU attachment to and dispersed prebound bacteria from the tubing, mimicking the phenotype of the ACICU Δabp1 Δabp2 strain (Fig. 5d, fig. S14, p=0.0052, one-way ANOVA).

### Clinical models of adhesin inhibitors

We next explored whether these adhesin inhibitors would function *in vivo* in clinical models of UTI. To test anti-FimH minibinder F7’s ability to combat uncomplicated UTI, mice were injected intraperitoneally with either 100 µg of the minibinder, mAb926, or a buffer-only control 24 hours prior to bacterial challenge. Mice were then challenged with model uropathogenic *E. coli* strain CFT073 from the clonal group ST73 (10^6^ CFU), and a second dose of protein was administered by intraperitoneal (IP) injection 24 hours after infection. We chose to administer the biologics 24 hours prior to and after challenge to maximize the potential *in vivo* effect. Forty-eight hours post-infection, bacterial burden in the bladder was quantified (Fig. 6a). F7-treated mice showed a significant (~2 log) reduction in bacterial bladder titers compared to the mock control group (p 0.0089, Mann-Whitney U test; Fig. 6b). This ~2 log reduction in bacterial bladder titers was similar to a control experiment comparing mAb926 treated mice to a mock control group (p 0.0001, Mann-Whitney U test; fig. S15).

**Fig. 6.**
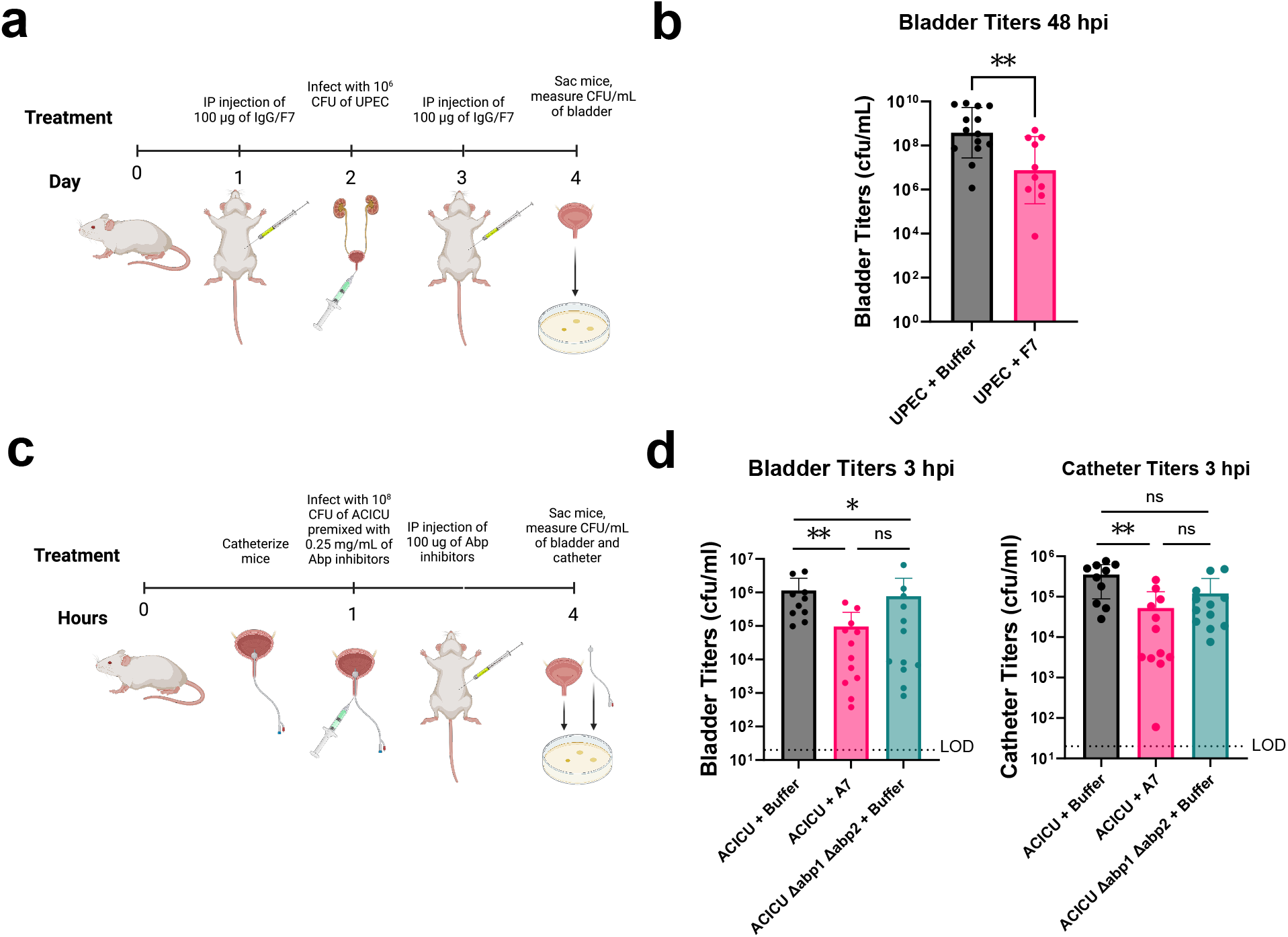
*In vivo* characterization of *de novo* designed adhesin inhibitors. **(A)** Schematic of UPEC UTI mouse model dosing regimen. Created with BioRender.com. **(B)** Titers of bacteria 48 hours post-infection (hpi) in the bladder after treatment with buffer (grey, n=14) are significantly higher (p=0.0089) than after treatment with minibinder F7 (pink, n=10). Data are presented as mean values +/− SD of n biological replicates. Mann-Whitney U test with two-tailed P value. ***P ≤ 0.001, **P ≤ 0.01, *P ≤ 0.05. Each data point represents one mouse; n indicates the number of mice per treatment group. Source data are provided as a Source Data file. **(C)** Schematic of ACICU CAUTI mouse model dosing regimen. Mice were infected with 10^8^ cells of ACICU in a CAUTI model. Their catheters and bladders were processed for counting CFUs 3 hpi. Created with BioRender.com. **(D)** Bladder (left) and catheter (right) titers of ACICU and ACICU Δabp1 Δabp2 3 hpi. ACICU were treated with buffer control (gray, n=9) or A7 (pink, n=12). ACICU Δabp1 Δabp2 were treated with buffer (teal, n=12). Bladder titer buffer vs A7 treatment p=0.0035. Bladder titer WT ACICU buffer vs ACICU Δabp1 Δabp2 buffer treatment groups p=0.0453. Catheter titer buffer vs A7 treatment p=0.0006. Data are presented as mean values +/− SD of n biological replicates. ne-way ANOVA with Tukey’s multiple comparisons test. **P ≤ 0.01, *P ≤ 0.05. Each data point represents one mouse; n indicates the number of mice per treatment group. Source data are provided as a Source Data file.

We next tested anti-Abp minibinder A7 in a mouse model of CAUTI. Mice were catheterized and immediately infected with *A. baumannii* ACICU (10 CFU) premixed with 0.25 mg/mL of minibinder or buffer. After 1 hour, mice were treated with IP injections of 100 µg of A7 or a buffer-only control. After 3 hours, bacterial burden in the bladder and catheter of each mouse was quantified (Fig. 6c). Mice that were treated with A7 exhibited significantly reduced bacterial loads in the bladder (p = 0.0035, one-way ANOVA) and catheter (p = 0.0006, one-way ANOVA) compared to mock-treated controls (Fig. 6d). This reduction in bacterial burden aligns with previous studies showing only modest (~1 log) attenuation of bacterial burden by ACICU Δabp1 Δabp2, suggesting alternative adhesion mechanisms may be involved in this mouse model [32]. This proof-of-concept experiment lays the ground for future antibiotic-sparing strategies against pathogens of great concern.

### Optimization of adhesin inhibitors

We explored two routes to generating higher affinity designs. First, we used ProteinMPNN to resample sequences based on the designed backbone of minibinder F7, and selected variants with improved AF2 Initial Guess scores; this led to the identification of a new minibinder (C8) exhibiting ~10-fold higher affinity for FimH (LAS K_d_=15.0nM; HAS K_d_=243nM) and more than ~30-fold greater inhibitory activity in an inhibition ELISA with an IC50 of 37 nM (95% CI: 31-47 nM; fig. S16, S17, table S2). Second, to enhance affinity via avidity effects, minibinder A7 was fused to *de novo* designed homo-oligomeric domains [37], generating multivalent binders. Oligomeric constructs such as oligomer C11 displayed significantly increased binding affinity to both adhesins, with K_d_s in the picomolar range for Abp1D and Abp2D (fig. S18). In cellulo, oligomer C11 exhibits a significantly lower inhibition ELISA IC50 (0.36 nM) than A7 (95% CI: 0.15-0.61 nM; fig. S19, table S2). Such affinity and avidity-increasing strategies should be generally applicable for further improving the *in vivo* efficacy of designed anti-adhesin minibinders.

## Discussion

In this study we show that *de novo* protein design tools can be employed to design potent and specific minibinder inhibitors of bacterial adhesins, virulence factors critical for colonization and biofilm formation in a broad range of bacterial infections, including UPEC UTI and *A. baumannii* CAUTI. The choice of target was key to *in vivo* success, as the extracellular nature of these adhesins made them readily accessible to the designed proteins. Being able to specifically target minibinders to the substrate binding epitope using RFdiffusion with hotspot conditioning led to neutralizing binders without the need for extensive antibody screening or knowledge of the native substrates. The conversion of FimH HAS to LAS by minibinder F7 shows that computational design allows not only the selection of the target epitope but also the target conformation. Targeting the substrate binding pocket of these adhesins should make it unlikely that resistance to these binders will arise quickly, as any mutation to the binding pocket may also disrupt adhesion. The specificity of these designed proteins for their targets should enable selective depletion of these pathogens without affecting commensal microbiota and limit pressure among the broader bacterial community to evolve resistance to the minibinders [38]. Future studies should explore whether resistance can be evolved to escape these minibinders and how they affect the broader microbiome. Although we show that these minibinders were specific to their target adhesin and do not bind their off-target adhesins, future studies will be needed to assess whether these minibinders exhibit off-target binding to other (i.e. host) proteins.

Future clinical development of the adhesion-neutralizing therapeutics demonstrated here will require characterization and possibly additional optimization of the cross-strain reactivity, binding specificity, route of administration, pharmacokinetics, and bioavailability of these designed proteins. Although some IP-delivered miniproteins are known to clear from serum quite quickly with half-lives less than 15 minutes, the half-lives for these specific miniproteins to filter through the kidney is unknown and will be determined in future work to optimize the dosing regimen [6]. Furthermore, we believe that fusions of these miniproteins to albumin binding domains or Fc-domains can be used to help tune the pharmacokinetic profiles of these anti-adhesin minibinders [39, 40]. Future head-to-head comparisons of the optimized minibinders with existing anti-adhesin drugs like mAb926 or mannoside drugs in *in vivo* studies will be helpful in identifying the strengths and limitations of this therapeutic modality. In the context of treating catheterized patients, another option for delivering minibinders is intravesical instillation through the urinary catheter directly into the bladder as has been implemented in previous clinical trials [41]. Future studies should also explore the utility of these anti-adhesin minibinders in treating more complex *in vivo* UTI models such as chronic infection models where existing treatments have limitations.

The therapeutic strategy of blocking adhesin activity using minibinders has potential utility against non-UTI bacterial pathogens such as *Staphylococcus aureus, Streptococcus pneumoniae;* and *Salmonella* [42, 43]. The strategies developed here should also be applicable to combat the adhesion and colonization of pathogens beyond bacteria, including fungi and amoebae [44, 45, 46].

## Materials and Methods

### Design

Ten thousand backbone designs were generated per target using the target crystal structure as input (PDB codes; Abp1d: 8DEZ, Abp2d: 8DF0, FimH HAS: 1UWF, FimH LAS: 3JWN) as well as hotspots (Abp1d: residues 55/93/110, Abp2d: 55/93/110, FimH: residues 54/133/137/51 or 1/13/48/138 or 52/55/135/140/164) corresponding to the substrate binding pocket residues. 20 sequences were assigned per backbone with ProteinMPNN (using default settings and temperature = 0.01) before AF2 Initial Guess filtering (using default settings and recycles = 3). Designs with pAE under 10 and pLDDT over 80 were then resampled using partial diffusion (using default settings and diffuser.T = 15), followed by MPNN and AF2 as before. This partial diffusion process was repeated once more. Abp backbones were then filtered to confirm their proximity (residues within 5 Å of the hotspots) to these “hotspot” residues and their likelihood of forming stable monomers, assessed by secondary structure topology analysis. Final designs were selected using AF2 Initial Guess metrics for the complex (pLDDT_binder ≥ 90, pAE_interaction ≤ 6.5), Rosetta interface metrics (DDG ≤ −30, SAP_score ≤ 40; using default settings and ddg scorefunction = “beta_nov16”), and AF2 monomer metrics for the minibinder alone (pLDDT_monomer ≥ 90).

### FimH expression, purification, and biotinylation

The lectin domain plasmid, containing a gene for ampicillin resistance and a His-tag, was transformed into *E. coli* BL21 cells and grown overnight at 37°C on LB agar plates supplemented with ampicillin (100 µg/mL). The resulting colonies were used to inoculate 1 L LB cultures supplemented with ampicillin, and the cultures were grown at 37°C with shaking at 200 rpm until reaching an optical density at 600 nm (OD_600_) of 0.6-0.7. The temperature was then reduced to 22°C for 1 hour prior to induction with 0.5 mM IPTG, followed by overnight incubation at 22°C. Cells were harvested by centrifugation at 7,000 × g for 15 minutes at 4°C. Periplasmic extraction was performed using osmotic shock. Specifically, the cell pellet was resuspended in 20 mM Tris-HCl, pH 8.0, containing 30% (w/v) sucrose (buffer B) and incubated at 4°C for 1 hour. Cells were pelleted by centrifugation at 18,000 × g and resuspended in 20 mM Tris-HCl, pH 8.0 (buffer C), and incubated at 4°C for 1 hour. The suspension was centrifuged at 33,000 × g for 15 minutes, and the resulting supernatant was collected as the periplasmic extract. The lysate was applied to a gravity-flow nickel affinity column and washed with 20 mM NaH_2_PO_4_, 300 mM NaCl, pH 8.0 (buffer D). Bound protein was eluted with the same buffer containing 500 mM imidazole. The eluate was further purified by size-exclusion chromatography using a Superdex 75 column equilibrated with 20 mM Na_2_HPO_4_, 100 mM NaCl, and 0.5 µM EDTA pH 6.0 (buffer E). The purified protein was concentrated using a 10 kDa molecular weight cutoff (MWCO) centrifugal filter at 4000 × g. Protein concentration was determined by measuring absorbance at 280 nm using a NanoDrop.

### Abp expression, purification, and biotinylation

The target receptor-binding domain proteins were expressed and purified from C600 *E. coli* with expression plasmids as described previously. [32] Briefly, protein was expressed by addition of 100 µm IPTG for 1 h and harvested by periplasmic extraction. RBD protein constructs and mutants were first purified by cobalt affinity chromatography, followed by cation (S) exchange chromatography. Purified protein was subsequently dialyzed or buffer exchanged into the appropriate buffer. Protein was then biotinylated using the EZ-Link™ NHS-PEG4 Biotinylation Kit (ThermoFisher, Cat#A39259) according to the manufacturer’s instructions.

### cDNA display

cDNA display was performed similarly to previously reported protocols for cDNA display [47] and Click Display [48], with major modifications. Note that the protocol used here was in development at the time of this project and may not represent the most efficient version of cDNA display, but it was sufficient for our work to identify binders. First, the library was amplified from Agilent oligo pool DNA using primers ACH_OLG_85_pETCON_F and ACH_OLG_86_pETCON_R and Kapa HiFi polymerase with 1× EvaGreen following the manufacturer’s instructions. A BioRAD C1000 Touch CFX96 Real-Time thermocycler was used to track amplification progress: hot-start activation at 95°C for 2 min; 39 cycles of 98°C for 20 s (denaturation), 65°C for 15 s (annealing), and 72°C for 45 s (extension); and a final extension at 72°C for 30 s. An initial test reaction determined the cycle at which amplification was complete, and a second scaled reaction (50 µL) was stopped at the cycle corresponding to half of the maximum fluorescence signal observed in the test run. The product size was then verified on a 1 % agarose gel, and the library was purified using SPRIselect Bead-Based Reagent before elution in nuclease-free water following the manufacturer’s instructions. For Golden Gate assembly, a 15 µL reaction containing the lCD9_1-3_inner (genbank file and sequence provided in Supplementary Data File 4) backbone (11 ng/µL) and the insert library (3.75 ng/µL) was incubated at 37°C for 1 h, followed by heat inactivation at 65°C for 5 min. Next, 0.75 µL of this Golden Gate reaction was added to a 50 µL Q5 PCR reaction (NEBNext Q5 Ultra II) using an annealing temperature of 65°C, and the mixture was amplified for 10 cycles. The final design library, which reintroduced the T7 promoter and display elements, was quantified by agarose gel and used without further purification in subsequent PUREfrex reactions. For *in vitro* transcription, translation, and mRNA-peptide conjugation, PUREfrex components were thawed on ice. In a low-binding PCR tube, a 25 µL PUREfrex reaction was assembled by combining 22.5 µL of PUREfrex Reaction Mix (including DnaK Mix used as a 20x stock), 2.5 µL of library DNA (approximately 15-25 ng), and 2 µM puromycin-modified cDNA oligo (which was prepared in the same manner as reported in click display.[48] The tube was gently mixed, spun briefly (≤ 1,000×g for 10 s), then incubated at 37°C for 20 minutes to allow coupled transcription and translation. Without cooling, the reaction was immediately exposed to 365 nm UV light (UVP UVL-28 EL Series UV Lamp, 95-0248-01) for 30 seconds to induce covalent linkage between the mRNA and the puromycin moiety. Next, 0.5 U/µL AcuI restriction enzyme and 1 µL GroEL/GroES chaperonin complex (GroE Mix used as a 20x stock) were added directly to the reaction, which was incubated at 37°C for 40 minutes and then at room temperature (~ 23°C) for 20 minutes to promote proper folding of displayed peptides. Ribosome dissociation was achieved by adding EDTA (pH 8.0) to a final concentration of 30 mM and incubating at room temperature for 5 minutes to chelate Mg2^+^. MgCl_2_ was then added to 30 mM to quench the remaining EDTA, followed by another 5-minute incubation at room temperature. Immediately thereafter, reverse transcription was initiated by diluting the reaction (42.5 µL) to 150 µL in SuperScript IV master mix (final concentrations of 1x SuperScript IV Buffer, 0.5 mM dNTPs, 10 mM DTT, and 2.66 U/µL SuperScript IV RT enzyme). The mixture was incubated at 50°C for 30 minutes. To purify the resulting cDNA-protein fusions via their N-terminal His-tags, the reverse transcription reaction was first diluted with 350 µL His Binding Buffer (10 mM Imidazole, 12 mM Phosphate, 500 mM Sodium Chloride, 2.7 mM Potassium Chloride, pH 7.4) and desalted using a 30 kDa cutoff Amicon spin concentrator and centrifuged at 14,000×g for 5 minutes at 4°C to reduce DTT concentration below 1 mM (for the His Mag Sepharose Beads). His Mag Sepharose Ni Beads were blocked with 100 µL of Starting Block buffer for 20 minutes before use. The retentate (~ 50 µL) was then incubated with blocked His Mag Sepharose Ni beads (50 µL bead slurry per 25 µL starting PUREfrex reaction) in His Binding Buffer for 1 minute. Beads were collected on a magnetic rack, washed four times with 100 µL Starting Block plus Tween and DNA buffer (SBTD, 1x Starting Block Buffer pH 7.4, 0.05 % Tween-20, 0.3 mg/mL sheared salmon sperm DNA) to remove unbound material, and then incubated in 100 µL His Elution Buffer (500 mM Imidazole, 12 mM Phosphate, 500 mM Sodium Chloride, 2.7 mM Potassium Chloride, pH 7.4) for 5 minutes at room temperature. The eluate containing purified cDNA-protein fusions was transferred to a fresh tube and subjected to three consecutive spin/dilute cycles (14,000×g, 15 minutes, 4°C) with 400 µL 1x PBS buffer pH 7.4 (137 mM NaCl, 2.7 mM KCl, 8 mM Na2HPO4, and 2 mM KH2PO4) each time to lower the imidazole concentration. For affinity panning, 10 µL Dynabeads Streptavidin T1 were transferred to a PCR tube and blocked with 100 µL Starting Block Buffer for 20 minutes on a rotator at room temperature. After placing the beads on a magnetic rack and discarding the supernatant, they were washed once with 100 µL SBTD buffer. The beads were then incubated with 100 µL SBTD containing 1000 nM biotinylated FimH (either the high-affinity wild-type (HAS) or the V27C/L34C low-affinity mutant (LAS)) for 20 minutes at room temperature to immobilize the target via streptavidin-biotin binding. After removing the unbound target and washing once with SBTD, the beads were resuspended in 100 µL SBTD containing the purified cDNA-protein fusions and incubated for 10 minutes at room temperature. Beads were washed four times with 100 µL SBTD on the magnetic separator, then transferred to a fresh tube after the final wash. To recover bound cDNA templates, beads were resuspended in 42 µL 10 mM Tris buffer pH 8. For downstream library regeneration, two parallel PCRs were performed on 2 µL of the panned bead slurry: one “regeneration” PCR to reintroduce T7 promoter and display elements (primers ACH_OLG_16_lCD_template_full_1_R and ACH_OLG_28_lCD_template_full_3_F), and one “NGS” PCR with primers containing Illumina overhangs (ACH_OLG_68 to ACH_OLG_75 in an equimolar mix). NGS primers contained a blend of differing length N bases to improve read quality.[49] Each 50 µL PCR contained 2 µL bead slurry, 25 µL NEBNext Q5 Ultra II 2× Master Mix, and 0.5 µM of each primer, with cycling conditions of 98°C for 30 seconds, 25 cycles of 98°C for 10 seconds/60°C for 20 seconds/72°C for 20 seconds, and a final extension at 72°C for 2 minutes. PCR products were verified on a 2 % agarose gel, gel-purified if necessary, and quantified by Qubit. The “Regeneration” amplicon was used directly in the next round of PUREfrex display without purification, while the “NGS” amplicon was submitted for high-throughput sequencing via Illumina MiSeq. Three total rounds of selection were performed, each with 1000 nM of either HAS or LAS antigens. Throughout the process, library recovery after antigen pull-down was monitored by qPCR but was not used for decision-making about when to stop performing selections. qPCR was performed on a BioRAD C1000 Touch CFX96 Real-Time thermocycler using the Luna® Universal Probe qPCR Master Mix from NEB following the manufacturer’s instructions. Primers ACH_OLG_9_lCD_qPCR_F, ACH_OLG_10_lCD_qPCR_R, and ACH_OLG_11_lCD_qPCR_P were used to quantify the total amount of recovered display product. 2 µL of the bead slurry containing recovered displayed proteins was diluted in 78 µL of 10 mM Tris buffer pH 8. For qPCR reactions, 4 µL of this dilution was used in a 10 µL qPCR reaction. Primers and reagents used in this section are listed in Supplementary Data File 5.

### Yeast surface display

For the yeast transformation, 50-60 ng of pETcon3, digested with NdeI and XhoI restriction enzymes, and 100 ng of the insert (PCR amplified from Twist oligonucleotide chip) were transformed into Saccharomyces cerevisiae EBY100 following the protocol previously described [2]. EBY100 cultures were cultivated in C-Trp-Ura medium with 2% (w/v) glucose (CTUG). To induce expression, yeast cells initially grown in CTUG were transferred to SGCAA medium with 0.2% (w/v) glucose and induced at 30°C for 16-24 h. During the “expression” sort, the induced yeast cells were washed with PBSF (PBS with 1% (w/v) BSA), stained for 30 minutes with a FITC-conjugated anti-C-Myc antibody (Immunology Consultants Laboratory, Cat. #CMYC-45F, 1:100 dilution) and PE-conjugated streptavidin (SAPE; Thermo Fisher, Cat. #S866, 1:100 dilution) in the absence of a biotinylated ligand, and the FITC-A+/PE-A-population was collected using a FACS sorter (Sony). The expression sort helps remove yeast cells that are not expressing any designed protein and cells that nonspecifically bind SAPE. For sorts 1 and 2, induced cells were stained with avidity (coincubation with SAPE) at room temperature for 30 minutes using 1 µM of biotinylated adhesin targets and FITC Anti-Myc mAb. Subsequently, cells were washed, and resuspended in PBSF, and single cells were individually sorted based on each target (FITC-A+/PE-A+) on a FACS sorter. For sort 3, cells were stained without avidity first at room temperature for 30 minutes using 10-fold titrations of biotinylated adhesin targets (from 100 nM to 1 nM), followed by subsequent washes in PBSF, and then stained for an additional 30 minutes at room temperature with SAPE and FITC Anti-C-Myc mAb. Again, the stained cells were washed, resuspended in PBSF, and individually sorted based on each target (FITC-A+/PE-A+) on a FACS sorter. Sorted cells were then grown to saturation in CTUG, harvested, and samples were prepared from them for next generation sequencing on a MiSeq (Illumina).

### Minibinder protein expression and purification

Minibinders were expressed in *E. coli* BL21 (NEB). Briefly, the DNA fragments encoding the design sequences were assembled into pET-29 vectors via BsaI golden gate assembly (NEB) and further transformed into BL21 strain with heat-shock. Protein expression was induced by the autoinduction system, and proteins were purified with immobilized metal affinity chromatography (IMAC). The elutions were further purified by FPLC SEC using Superdex 75 10/300 GL or Superdex 10/300 200 columns (GE Healthcare). Protein concentrations were determined by NanoDrop (Thermo Scientific) and normalized by extinction coefficients. Proteins were diluted to the appropriate concentrations in 1x PBS (for cell assays) or 1X HBS-EP+ buffer (for binding assays). Endotoxin was removed during protein production for the *in vivo* studies, following previously described methods [37].

### Circular dichroism and thermal melts

The secondary structure content was evaluated by CD in a Jasco J-1500 CD spectrometer coupled to a Peltier system (EXOS) for temperature control. The experiments were performed on quartz cells with an optical path of 0.1 cm, covering a wavelength range of 200-260 nm. The CD signal was reported as molar ellipticity (MRE). The thermal unfolding experiments were followed by a change in the ellipticity signal at 222 nm as a function of temperature. Proteins were denatured by heating at 1 °C per minute from 20 to 95 °C.

### SPR binding assays

SPR experiments were conducted using a Biacore™ 8K instrument (Cytiva) and analyzed with the accompanying evaluation software. Biotinylated adhesins were immobilized on a streptavidin sensor chip (Cytiva) at a concentration of 0.4 µg/ml for Abp1D/Abp2D and 1.2 µg/ml for high-affinity state wild-type FimH and a low-affinity state mutant (L34K). Increasing concentrations of protein binders were flown over the chip in 1X HBS-EP+ buffer (Cytiva). Final fits were performed with Biacore™ Insight Software (Cytiva). FimH HAS fits were performed using the two-state reaction model. All other fits were done using the 1:1 binding model.

### Crystallization sample prep and analysis

Crystallization experiments for the binder complex were conducted using the sitting drop (for F7-FimH) or hanging drop (for A7-Abp2D) vapor diffusion method. Crystallization trials were set up in 200 nL drops using a 96-well format by Mosquito LCP from SPT Labtech. Crystal drops were imaged using the UVEX crystal plate hotel system by JANSi. For F7-FimH, diffraction quality crystals appeared in 0.2 M Sodium malonate, pH 7.0, 20% w/v Polyethylene glycol 3,350. Crystals were flash-cooled in liquid nitrogen before shipping to the synchrotron for diffraction experiments. Diffraction data were collected at the NSLS2 beamline FMX (17-ID-2). For A7-Abp2D, crystals appeared in 0.1M Sodium acetate, pH 4.6, 0.15 M Ammonium acetate, and 25% PEG4000. Diffraction data were collected at the ALS 4.2.2 beamline. X-ray intensities and data reduction were evaluated and integrated by XDS and merged/scaled by Pointless/Aimless in the CCP4i2 program suite. Structure determined by molecular replacement using a designed model using Phaser. Following molecular replacement, the model was improved and refined by Phenix. Model building was performed by COOT in between refinement cycles.

### NMR Sample Preparation, Data Collection, and Analysis

^15^N-labeled FimH lectin domain (LD) was expressed in *E. coli* grown in M9 minimal media containing ^15^N-ammonium chloride as the sole nitrogen source. Protein expression was induced with 1 mM IPTG. Samples were prepared in NMR buffer containing 20 mM sodium phosphate (pH 6.0), 0.5 µM EDTA, and 100 mM NaCl. The LD was prepared at a final concentration of 100 µM. To generate protein-ligand complexes, a 7-fold molar excess of the F7 minibinder was added to wild-type LD, and a 5-fold excess was added to L34K-LD. ^15^N-HSQC spectra were acquired at 298 K on a Bruker AVANCE 800 MHz spectrometer equipped with a cryoprobe. Spectra were processed using NMRPipe and analyzed with NMRViewJ [50, 51].

### Acinetobacter Abp2D Monoclonal Antibody Generation

All animal work was conducted with the approval of the Washington University Institutional Animal Care and Use Committee. Five 6-8 week old C57BL/6 mice (Jackson laboratories) received 2 doses 3 weeks apart of 50 µg of Abp2D mixed 1:1 by volume with a squalene oil-in-water adjuvant (Addavax, Invigen). Mice were housed in standard Washington University barrier animal housing with a 12/12 light dark cycle at 22°C and 30% humidity. 5 days after the second dose, mice were sacrificed and inguinal and iliac lymph nodes were collected into RPMI containing 2% fetal bovine serum and manually homogenized using the back of a syringe plunger. Cells were filtered through 75 µm mesh, washed 1x, and counted. All washes for the staining process were performed in PBS containing 2% fetal bovine serum and 2 mM EDTA. All cells for each sample were stained with the following panel: live/dead Zombie Aqua, CD3 APC-Cy7, CD19 PE, Fas APC, IgD FITC, CD38 PE-Cy7, CD138 Pacific Blue. CD3-CD19+/IgDlo Fas+/CD38-CD138+ cells were single-cell sorted on a BD FACSAria at the Washington University Flow Cytometry Core into 96-well plates. Lymph nodes from naïve mice were used to set gates. 2 plates of B cells were sorted on 08-21-20. A second B cell sort of 2 plates was conducted on 01-23-21. For the 2nd B cell sort, the flow panel used was CD4-CD19+/IgDlo/B220-CD138+. Mabs were cloned from sorted B cells as described previously. [52, 53] Briefly, B cell RNA was converted to cDNA via RT-PCR. cDNA was used as template for 4 sets of nested PCR, one each with primers against IgG, IgM/IgA, Ig*κ* and Ig*λ* variable regions. Wells with successful amplification from heavy and light chains were sequenced (Azenta) and analyzed using the IMGT V-quest web tool (https://www.imgt.org/IMGT_vquest/input) to isolate clonally distinct mAbs. Heavy and light chain plasmids for each clone were co-transfected into Expi293 cells (Thermo Fisher, Cat. #A14527) using the Expifectamine transfection reagent according to the manufacturer’s instructions. Antibodies were purified on protein A agarose. To determine Abp2D antigen specific mAbs, ELISA plates were coated with 1 µg/ml Abp2D, 0.1 µg/ml bovine serum albumin (as a negative control) or 0.1 µg/ml anti-Ig antibody (as a positive control for mAb expression) overnight at 4°C. The next morning, plates were washed 1x with PBS-T (1x PBS+ 0.05% Tween-20) and blocked with P10 (PBS + 10% fetal bovine serum) for 1.5 hours at room temperature. Supernatants were diluted 1:30 in P10 and incubated 1 hour at room temperature. After washing 3x with PBS-T, the secondary antibody (anti-human IgG-HRP, Jackson ImmunoResearch, Cat. #109-035-088) was diluted 1:2,500 in P10 and incubated 1 hour at room temp in the dark. Plates were washed 3x with PBS-T followed by 3x with PBS. Detection reagent (10 ml phosphate-citrate buffer, 4 mg o-Phenylenediamine dihydrochloride, and 33 µl 3% H2O2 per plate) was added to the plates, incubated 5 minutes, and quenched with 1 M HCl. Absorbance (490 nM) was quantified using the BioTek ELx800 plate reader on the OD490 setting.

### Fibrinogen ELISA with purified adhesins

Flat-bottom microplates (Greiner Bio-One) were coated overnight at 4°C with human fibrinogen from plasma (150 µg/ml) (Sigma). The plates were blocked for an hour with 1.5% BSA in PBS, followed by washing with PBS (three times for 5 min each). Biotinylated adhesins at 500 nM were premixed with varying concentrations of Abp inhibitors (Fabs or minibinders), added to the coated wells, incubated 1 hour at room temperature or overnight at 4°C, and then washed three times to remove the unbound adhesins. After the washes, anti-streptavidin-HRP (BD Pharmagen, Cat# 554066, 1:5000 dilution) was allowed to bind for 1 hour before washing 3 times with PBS-T and development with TMB substrate (BD Pharmagen, Cat# 555214). After reactions were quenched with 1M H_2_SO_4_, absorbance at 450 nm was recorded.

### Fibrinogen bacterial ELISA assays

Flat-bottom microplates (Greiner Bio-One) were coated overnight at 4°C with human fibrinogen from plasma (Fisher Scientific, Cat. #NC9464127, 150 ug/mL). The plates were blocked for an hour with 1.5% BSA in PBS, followed by washing with PBS (three times for 5 min each). Bacterial strains were grown statically overnight in LB broth and washed and resuspended in PBS. For “inhibition” assays, a total of 100 µL of bacteria (final OD600 of 0.5) were premixed with varying concentrations of Abp inhibitors, added to the coated wells, and incubated for 2 hours at 37 °C, followed by PBS washes to remove the unbound bacteria. For “detachment” assays, 100 µL of bacteria (final OD600 of 0.5) were directly added to the fibrinogen-coated wells, incubated for 2 hours at 37°C, and then the wells were washed with PBS. Abp inhibitors were then added and allowed to incubate for 2 hours at 37°C, after which wells were washed with PBS to remove the unbound bacteria. Next, bacterial cells were fixed with formalin for 20 min at room temperature, followed by three washes with PBS containing 0.05% Tween 20 (PBS-T). Then, the plates were blocked overnight at 4°C with 1.5% BSA-PBS, followed by three washes with PBS-T. After the washes, the plates were incubated for 2 hours at room temperature with rabbit anti-Acinetobacter antisera (from Di Venanzio et al., 1:1,000). Plates were washed with PBS-T, incubated with anti-rabbit-HRP (KPL, Cat. #5220-0336, 1:1000 dilution) for 1 hour. After washing 3 times with PBS-T and development with TMB substrate (BD Pharmagen, Cat. # 555214), reactions were quenched with 1M H2SO4 and absorbance at 450 nm was recorded.

### Bacterial strains expressing FimH

The recombinant *Escherichia coli* K12 strain (AAEC191A) carrying pPKL114 plasmid containing the entire *fim* gene cluster from the *E. coli* strain K12 (but with the inactivated *fimH* gene) and pGB2-24 plasmid carrying *fimH* from the model uropathogenic E. coli strain J96 (resulting in *E. coli* KB23) or *Klebsiella pneumoniae* strain (resulting in *E. coli* cas665) were described previously [54, 55]. Clinical *E. coli* isolates from ST73 (strain 65-1H), ST95 (51-6B) and ST69 (45-3G), as well as strains from the global pandemic multi-drug resistant clonal groups ST131-H30 (38-9H) and ST1193 (53-6E) obtained from urine of patients with suspected UTI submitted for analysis to the clinical microbiology laboratory of the Kaiser Permanente Washington, Seattle [56]. The clonal identity was determined by fimC:fimH Sequence Typing [57]. To induce the type 1 fimbriae expression, bacteria were passed 3 times by overnight subculturing in Luria-Bertani broth without shaking. Before testing, bacteria were washed two times in PBS and adjusted for the desired OD_540_ in 0.1% BSA/PBS.

### Red blood cell agglutination inhibition and dispersion assays

Guinea pig red blood cells (RBC) were purchased from Colorado Serum Co (Denver, CO), were washed with PBS three times and diluted to 1% v/v. To normalize for the bacterial population expressing type 1 fimbriae, the washed bacteria were tested first for the working concentration to induce the RBC agglutination. Flat-bottom 96 well plates were quenched with 1% BSA-PBS, 30 µL of the 1:1 dilutions of bacteria (starting from OD=2) was placed in each well, mixed with 30 µL of RBC, the plate was placed into a shaker (Microplate Genie™; Scientific Industries, Bohemia, NY) and subjected to shaking at 600 rpm for 20 minutes. The plate was taken out, gently tapped on the corners to homogenize the well content for recording. The second highest dilution that induced a clear agglutination of RBC was chosen for the inhibition assays (for the recombinant bacteria, the working concentration was 4 x 10 CFU/mL). The working concentration of bacteria was mixed 1:1 with the minibinder, mannose or monoclonal antibody solution at the indicated concentration, prepared in 0.1% BSA/PBS, and tested for the RBC agglutination as described. For the RBC agglutination dispersion assay, the RBCs were pre-agglutinated with the working concentration of bacteria as described above. After the RBCs were pre-agglutinated, 15 µL of different concentrations of the minibinders or monoclonal antibodies were added on the top of the RBC-bacteria suspension followed by immediately re-shaking for 10 min. The final concentration of inhibitors used for this detachment assay was 5 µM for the minibinders and 5 nM for mAb926. MICs were calculated by visual inspection to determine the highest concentration where the RBC carpet was not fully formed (partially collapsed).

### Inhibition of *E. coli* adhesion

Microtiter 96-well plates were coated with 20 µg/ml of bovine RNAse B (Sigma-Aldrich) in 0.02 M NaHCO3 buffer at pH 9.6. The wells were quenched with 0.2% bovine serum albumin (BSA, Sigma-Aldrich) in PBS for 20 min. Bacteria expressing FimH-J96 (OD = 1) were mixed with different concentrations of minibinders for 30 min at 37°C and then allowed to adhere to the coated surface for another 1 h. After an extensive washing with PBS, plates were dried and bound bacteria were stained with 0.1% crystal violet (Becton Dickinson) for 20 min at room temperature (RT). The wells were washed with water and 50% ethanol was added to the wells. The absorbance was measured at 600 nm.

### Detachment of *E. coli* adhesion

Microtiter 96-well plates were coated with 20 µg/ml of yeast mannan (Sigma-Aldrich) in 0.02 M NaHCO3 buffer at pH 9.6. The wells were quenched with 0.2% bovine serum albumin (BSA, Sigma-Aldrich) in PBS for 20 min. Bacteria expressing FimH-J96 (OD = 1) were added into the wells and spun down at 4,800 × g for 5 minutes. After washing with PBS from unbound bacteria, 100 µL of 0.1% BSA/PBS was added with different concentrations of minibinders or antibodies. The plate was incubated for 1 hour at 37°C without shaking, followed by washes with PBS and crystal violet staining as described above.

### *E. coli* bladder cell adhesion assays

Bacterial cell suspensions (50 µL of 10^9^ CFU/mL) were mixed 1:1 with McCoy’s 5A medium with 2% heat inactivated fetal bovine serum, without added inhibitors, with 0.5% *α*-methyl-D-mannopyranoside, 5 µM of the minibinders, or with 5 nM of mAb926 and were added to monolayers of T24 human bladder epithelial carcinoma cells grown in 12-well plates. Bacteria were allowed to interact with the cells for 2 h at 37°C in 5% CO2. The cells were washed six times with PBS, fixed with cold 100% methanol for 1 min, air dried and stained with eosin Y followed by Hema3, both dyes diluted 1:2 with water. Representative microscopy images were taken at 20X magnification. The number of bacterial cells per T24 bladder cell were counted for at least 3 replicate images per treatment condition. T24 cells were obtained from ATCC (HTB-4). No authentication procedure and mycoplasma testing were performed.

### ACICU Biofilm assay

Overnight static cultures were normalized to an OD600 of 0.01in Lennox LB media (Fisher) and inoculated into sterile, round-bottomed, PVC 96-well plates (Corning Inc.) with 200 µL final volume per well. For biofilm inhibition, treatment or control proteins in PBS were added before growth. Plates were incubated for 24 hours under static conditions at 37°C with humidity to prevent evaporation. For measuring detachment, bacteria were allowed to form biofilms for 24 hours and then were treated with minibinder or control protein for 2 hours at 37°C. Unbound cells in each well were washed 3 times with water. The adhered biofilm was stained with 300 µL of 0.5% crystal violet solution dissolved in water for 15 min. After staining, wells were washed 3 times with water and left to completely dry. Crystal violet staining was dissolved in 200 µL of 30% acetic acid, and absorbance was measured at 600 nm.

### Dispersion of biofilm from fibrinogen-coated catheters

For immunohistochemistry quantification of bacterial and fibrinogen deposition on catheters, we used a modified protocol from the methods outlined in Andersen et al. and Di Venanzio et al. (PMCIDS: PMC8986317; PMC6591400). [36, 58] Silicon tubing (4 to 5 mm diameter; used in mouse catheter experiments) was cut into approximately 0.5 cm catheter sections and UV sterilized. Fibrinogen (Fisher Scientific, Cat. #NC9464127, 150 ug/mL) was allowed to coat the catheter sections overnight at 37°C. Catheters were then fixed in 10% formalin for 20 minutes at room temperature and washed 3x in PBS. Bacteria (OD = 0.5 in PBS) were added to catheters and allowed to bind at 37°C. For inhibition studies, bacteria and 100 nM of BSA or minibinder A7 were added simultaneously to the catheter sections and allowed to bind for 2 hours. For detachment studies, bacteria were first allowed to bind to the catheter for 1 hour, and then incubated for 1 hour with 100 nM of BSA or A7. Catheter sections were then washed 3x with PBS, fixed in 10% formalin for 20 min at room temperature, washed again 3x in PBS, and allowed to block overnight in blocking buffer (PBS solution with 1.5% BSA and 0.1% sodium azide). Primary antibodies (1:500 goat anti-fibrinogen (Sigma-Aldrich Cat# F8512, RRID:AB_259765), and 1:1000 rabbit anti-acinetobacter from Di Venanzio et al.) in blocking buffer were allowed to bind for 1 hour at room temperature. [36] After washing 3x in wash buffer (PBS with 0.05% Tween-20), secondary antibodies (1:10,000 LI-COR donkey anti-goat IRDye 800CW (LI-COR Biosciences Cat# 926-32214, RRID:AB_621846) and 1:10,000 LI-COR donkey anti-rabbit IRdye 680LT (LICOR Biosciences Cat# 926-68023, RRID:AB_10706167) in blocking buffer were allowed to bind for 1 hour, followed by 3x washes. Catheters were allowed to dry overnight at 4°C and imaged on an Odyssey Imaging System (LI-COR Biosciences). Experiments were performed in four independent replicates. The catheters shown in the Fig. 5d are from one representative replicate and were imaged together to ensure identical detection settings. For quantification in Fig. 5c, bacterial signal (red) on catheters was normalized by fibrinogen signal (green) using ImageJ [59] to account for variation in catheter size and fibrinogen coating.

### *In vivo* UTI mouse model with UPEC

Eight week old female BALB/c, mice (Charles River Laboratories) from the same cage were randomly sorted into two groups for each experiment. The first experiment compared the bladder titers of 21 mAb926-treated mice to 24 buffer-treated mice. The second compared the bladder titers of 14 buffer-treated mice to 10 F7-treated mice. For both experiments, mice were passively immunized with 100 µg IP injections 24 hours before challenge with 10^6^ CFU of *E. coli* CFT073. A second 100 µg dose of F7/IgG was administered IP 24 hours post-infection. The mice were euthanized 24 hours later (48 hours post-infection), after which their bladders were collected for bacterial load determination. The bacterial load was determined in units of CFUs/g of organ tissue. This study was approved and performed in accordance with the guidelines set by Harvard University under IACUC protocol # IS00000235-6 on April 24, 2023.

### *In vivo* CAUTI mouse model with *A. baumannii* ACICU

Six-to eight-week-old female C57BL/6 mice (Charles River Laboratories) were infected in a catheter-associated UTI model as previously described [32, 60]. Briefly, a 4-to 5-mm piece of silicone tubing (catheter) was placed in the bladder via transurethral insertion. Bacterial strains were grown twice overnight at 37 °C and resuspended in 1x PBS. Mice were infected immediately following implant placement with ~2 × 10^8^CFUs of bacteria. Where indicated, bacteria were premixed with 0.25 mg/mL of inhibitors (final concentration) in 50 µL total volume via transurethral inoculation. Mice were then immediately dosed with 100 µg of Abp inhibitors via intraperitoneal (IP) injection. At 3 hours post infection (HPI), mice were sacrificed, and bladders and implants were aseptically removed. The bacterial load present in each tissue was determined by homogenizing each organ in PBS and plating serial dilutions on LB agar supplemented with antibiotics when appropriate. Mice were housed in standard Washington University barrier animal housing with a 12/12 light dark cycle at 22°C and 30% humidity. All CAUTI studies were approved and performed in accordance with the guidelines set by the Committee for Animal Studies at Washington University School of Medicine under IACUC protocols 24-0279 (IACUC Protocol Approval Animal Welfare Assurance # D16-00245).

### Statistics & Reproducibility

Statistical analyses were performed using GraphPad Prism (v11). Experiments were repeated using independent biological replicates to ensure reproducibility. For *in vitro* experiments, a minimum of three independent biological replicates was performed unless otherwise indicated in the figure legends. For mouse infection experiments, sample sizes were based on prior studies using established UPEC and *A. baumannii* colonization models. No statistical method was used to predetermine sample size. No data were excluded from the analyses except for in the case of the CAUTI mouse model experiment in which a catheter was not found present in the mouse at 3 hpi. Mice were randomly assigned to treatment groups. The investigators were not blinded to allocation during experiments and outcome assessment.

## Data availability

The crystal structures generated in this study have been deposited in the Protein Data Bank (PDB) database under accession codes 9Q1V (FimH-F7) [https://doi.org/10.2210/pdb9Q1V/pdb] and 9Q1H (Abp2D-A7) [https://doi.org/10.2210/pdb9Q1H/pdb]. The FimH cDNA panning NGS sequencing data generated in this study have been deposited in the NCBI Sequence Read Archive (SRA) database under the BioProject ID PRJNA1508338 [http://www.ncbi.nlm.nih.gov/bioproject/1508338] and BioSample accession codes: SAMN62230579-SAMN62230585. All other data are available in the main text or the supplementary materials. Source data are provided with this paper.

## Acknowledgments

Crystallographic diffraction data were collected at the Northeastern Collaborative Access Team beamlines at the Advanced Photon Source, which are funded by the National Institute of General Medical Sciences from the National Institutes of Health (P30 GM124165) and the ALS 4.2.2 beamline (P30 GM124169). This research used resources of the Advanced Photon Source, a US Department of Energy (DOE) Office of Science User Facility operated by Argonne National Laboratory under contract no. DE-AC02-06CH11357. This research used resources (FMX) of the National Synchrotron Light Source II, a U.S. Department of Energy (DOE) Office of Science

User Facility operated for the DOE Office of Science by Brookhaven National Laboratory under Contract No. DE-SC0012704. The Center for BioMolecular Structure (CBMS) is primarily supported by the National Institutes of Health, National Institute of General Medical Sciences (NIGMS) through a Center Core P30 Grant (P30GM133893), and by the DOE Office of Biological and Environmental Research (KP1607011). Figures were made using Biorender.com. We would like to acknowledge Jacob M. Gershon, Hojae Choi, Yensi Flores Bueno, David S. Lee, Jeremiah Sims, Brian Coventry, Alex Shida, Jesús Bazán Villicaña,, and Jay Nix for materials, helpful discussions, and support; and L. Goldschmidt and K. VanWormer for general operations.

## Author contributions

Conceptualization: ACG, TRT, EDBL

Methodology: ACG, TRT, EDBL, PM, REK, PWE, KWD

Investigation: ACG, TRT, PM, DAS, EDBL, MC, AV, SB, DW, AH, KOT, AK, EJ, AKB, AM, PA, VT, IB, MRT, JSP, GI, AJS

Visualization: ACG, TRT, EDBL, PM, KLH

Writing: ACG, TRT, EDBL, KWD, KLH, DB, SJH, EVS

Supervision: DB, SJH, EVS, AHE

## Supplementary Materials

Figs. S1 to S19

Tables S1 to S5

Supplementary data files 1 - 5

