## Supplementary information for "*De Novo* Design of Miniprotein Inhibitors of Bacterial Adhesins"

**This PDF file includes:**

- Figs. S1 to S19
- Tables S1 to S4

### Supplementary Figures

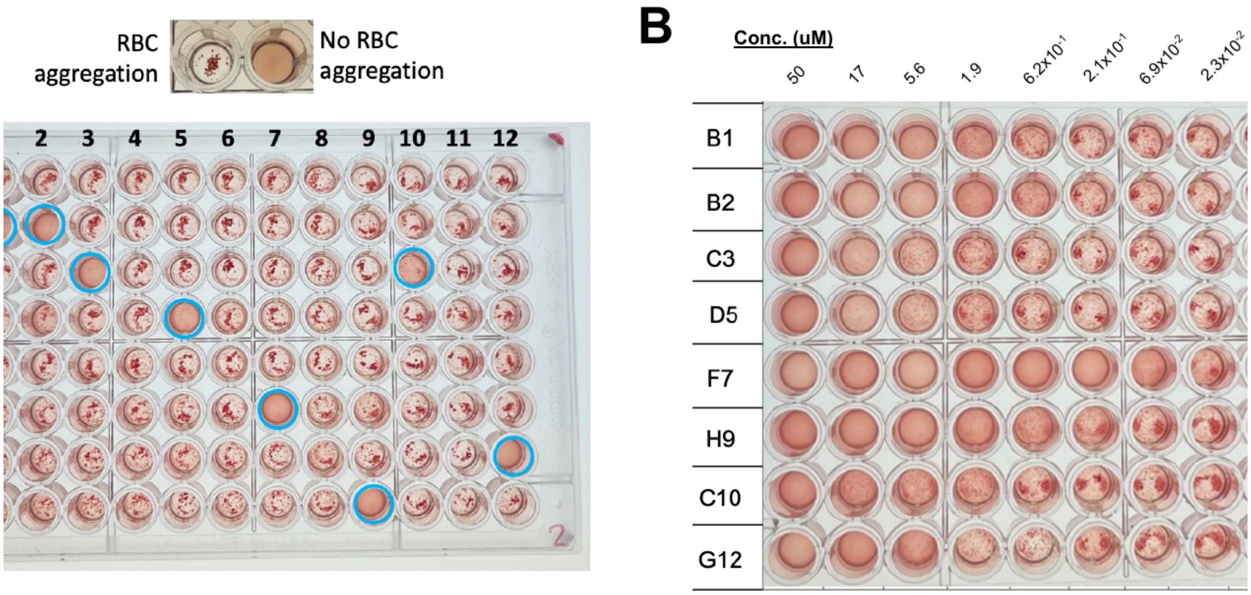


**Fig. S1. Initial RBC agglutination screen of designed FimH minibinders. (A)** RBC agglutination assay results for all enriched minibinders from cDNA display screening at a fixed concentration. Blue circles indicate binders that inhibit RBC agglutination. **(B)** Promising designs were then titrated at the indicated concentrations (shown in micromolar) to determine their RBC agglutination inhibition IC50 values.

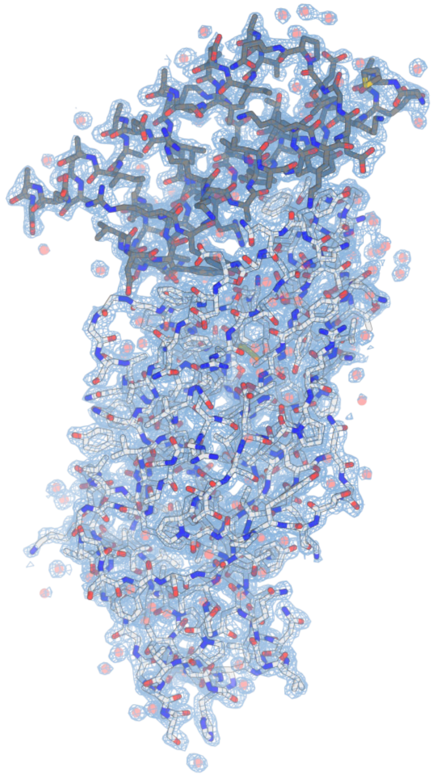

**Fig. S2. Electron density map of F7-FimH co-crystal.** 2mFo-DFc electron density maps contoured at 1 $\sigma$ shown in blue. F7 is shown in grey and FimH is shown in white.

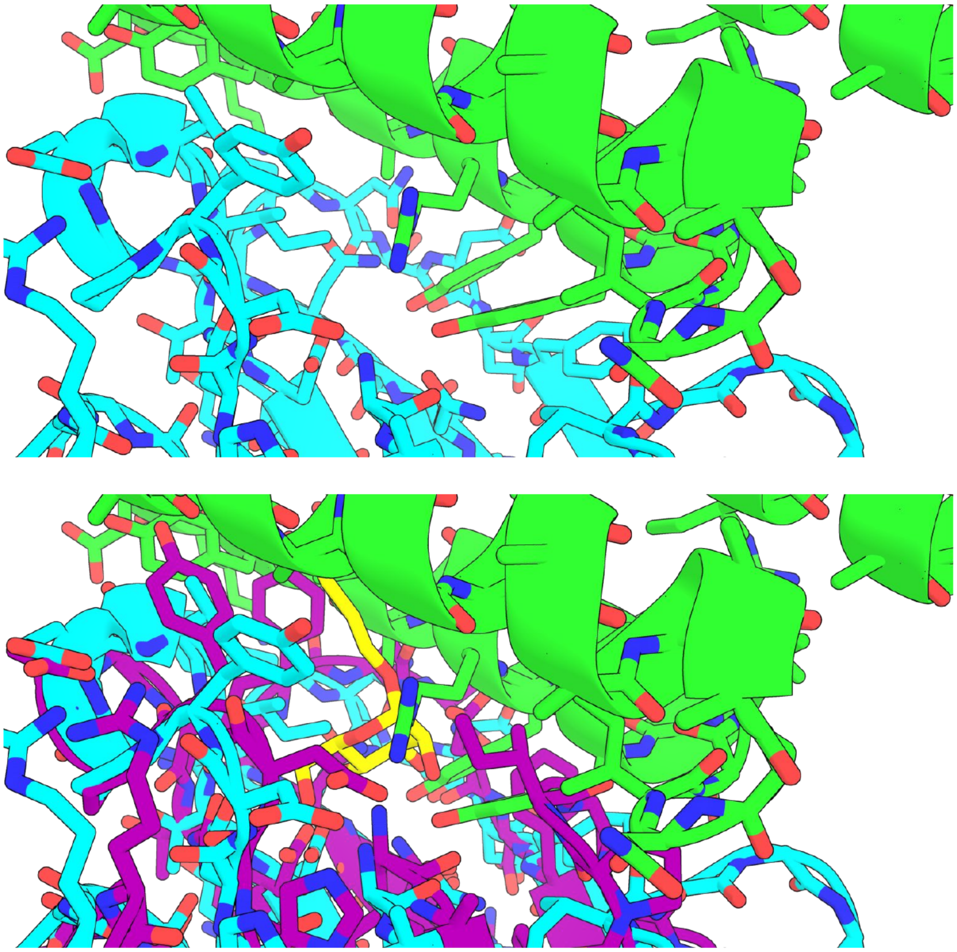

**Fig. S3. Closeup of the F7-FimH atomic interface. (A)** Closeup of the experimentally determined structure of F7 (green) bound to FimH (cyan). **(B)** Overlay of minibinder F7-FimH complex crystal structure with the structure of FimH (purple) bound to mannose (yellow; PDB code: 1UWF).

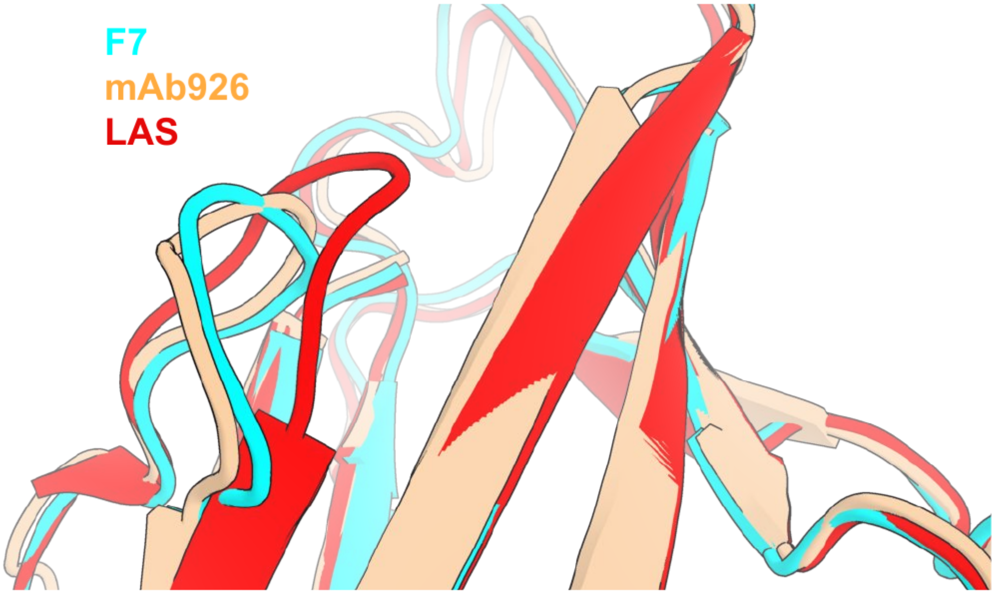


**Fig. S4. Cartoon overlay of FimH clamp loop.** The FimH clamp loop is shown in the F7-FimH complex (cyan), mAb926-FimH complex (PDB code 9ME5; wheat), or apo-LAS (PDB code 3JWN; red).

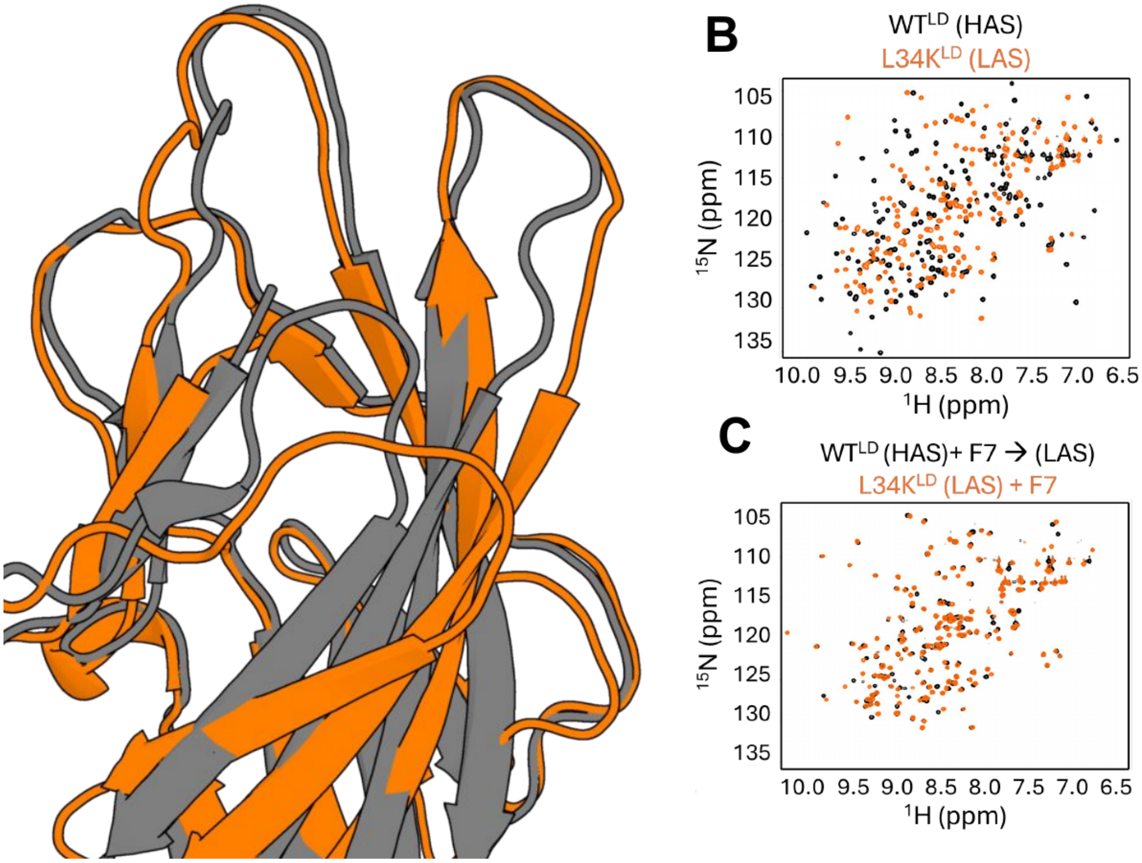


**Fig. S5. Minibinder F7 shifts the FimH conformational equilibrium. (A)** Overlay of FimH LAS (orange) and HAS (grey). **(B)** $^{15}$N-HSQC NMR spectra of FimH HAS (black) and LAS (orange) in the absence of any inhibitor. Note the general lack of overlap between the two spectra, showing pronounced differences between the two states. **(C)** $^{15}$N-HSQC NMR spectra of FimH HAS (black) and LAS (orange) in the presence of minibinder F7. Note the overlap induced by addition of F7, showing conversion of the HAS to the LAS. Source data are provided as a Source Data file.

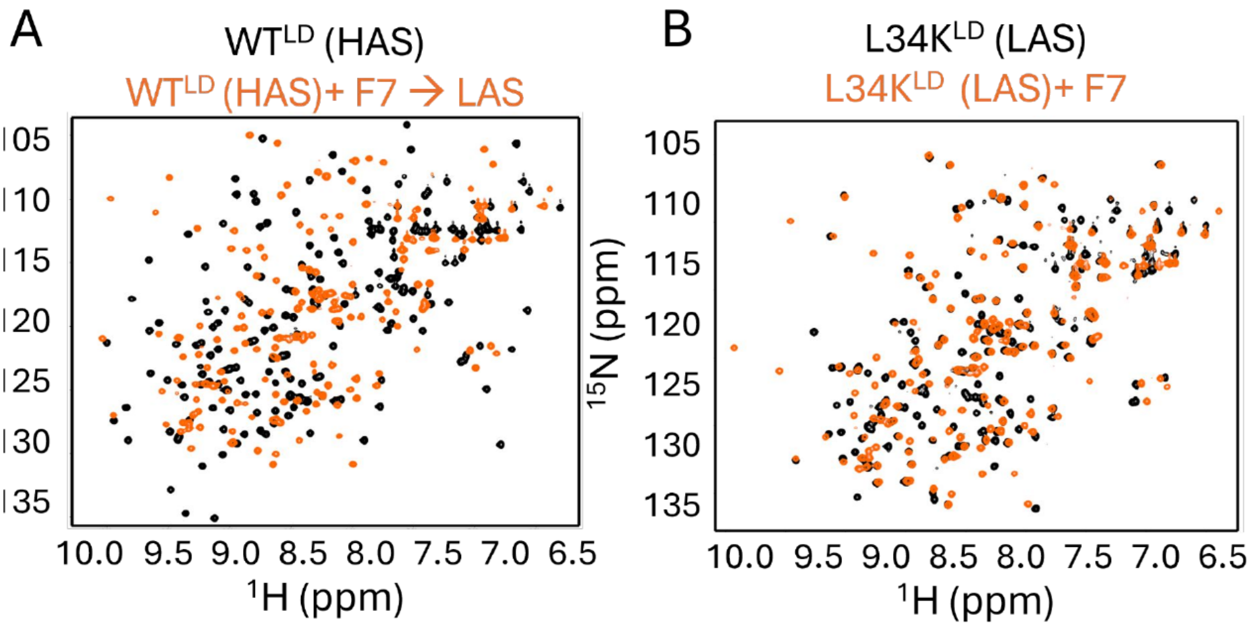


**Fig. S6. Overlay of FimH NMR spectra in presence and absence of minibinder F7. (A)** $^{15}$N-HSQC NMR spectra of FimH HAS (black) and HAS in the presence of minibinder F7 (orange). Note that the spectra shift markedly upon addition of F7. **(B)** $^{15}$N-HSQC NMR spectra of FimH LAS (black) and LAS in the presence of minibinder F7 (orange). Note that the spectra remain largely unchanged upon addition of F7. Source data are provided as a Source Data file.

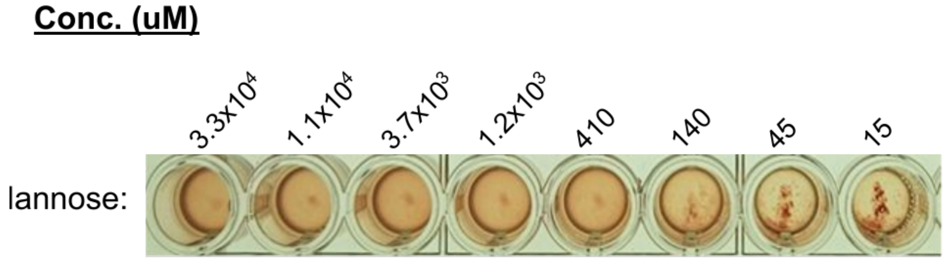


**Fig. S7. RBC agglutination inhibition by mannose.** The minimum inhibitory concentration of mannose on RBC agglutination is 140 µM, several thousand fold higher than F7 (69 nM).

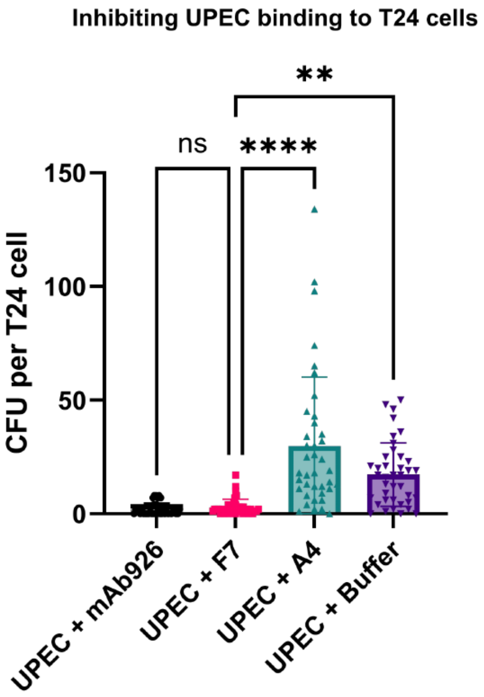


**Fig. S8. Quantification of UPEC adhesion to T24 bladder epithelial cells.** Addition of F7 (pink; mean = 2.85, SEM+/-0.57) and mAb926 (black; 2.0, SEM+/-0.42; F7-mAb926: n.s., ANOVA) significantly inhibits adhesion of UPEC to T24 cells relative to a buffer treatment (purple; mean = 17.26, SEM+/-2.24; F7-buffer: p<0.0012) and noninhibitory minibinder A4 control (teal; mean = 29.79, SEM+/-4.86; F7-A4: p<0.0001, ANOVA). Data are presented as mean values +/- SD. One-way ANOVA with Tukey's multiple comparisons test. Source data are provided as a Source Data file.

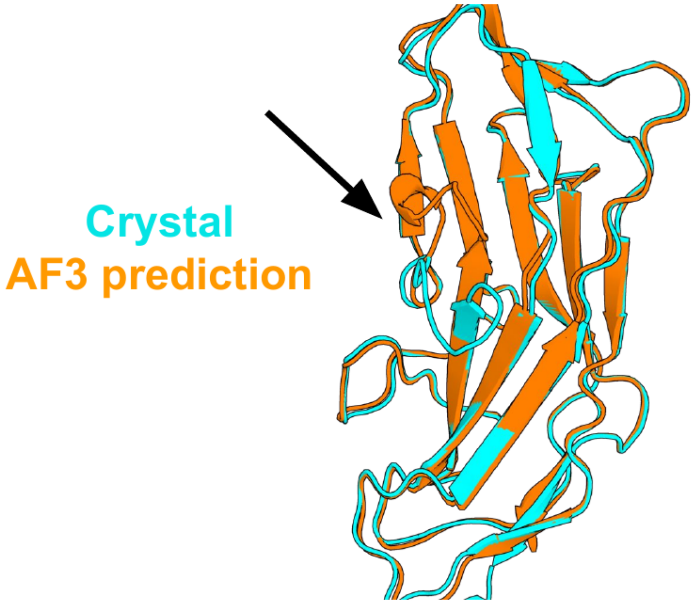


**Fig. S9. Cartoon overlay between the crystal structure of Abp2D and its AlphaFold2 model.** The arrow identifies the flexible anterior binding loop of Abp2D that is part of its putative fibrinogen binding pocket.

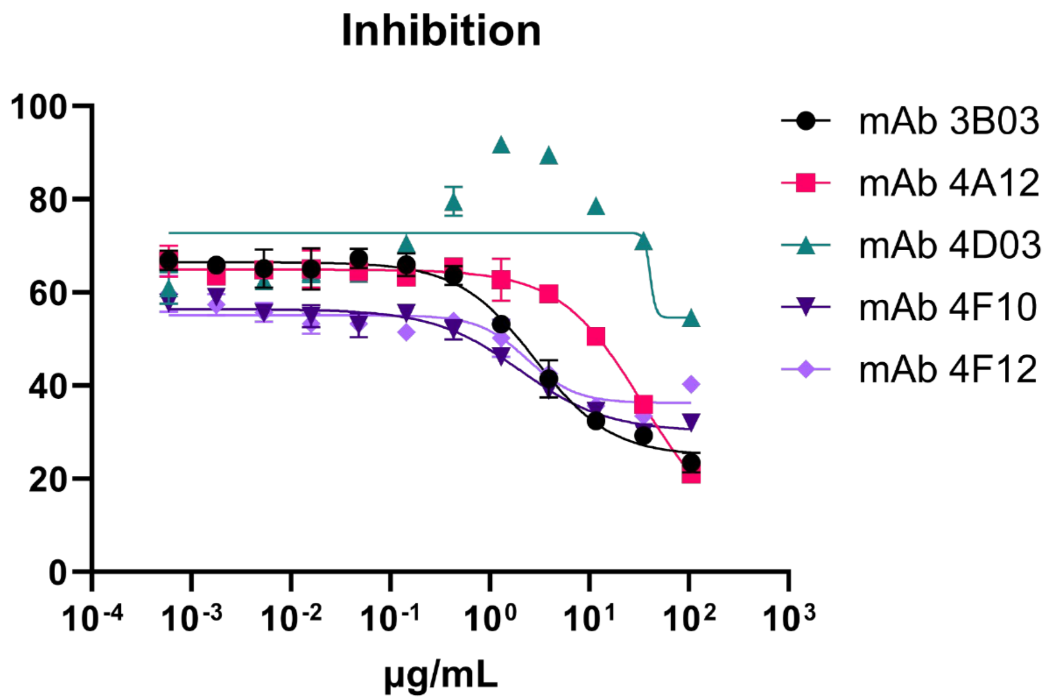


**Fig. S10. Purified adhesin ELISA for noninhibitory Abp2D mAbs.** ELISA results for mAb-mediated inhibition of purified Abp2D binding to fibrinogen. The data provided contain 3 replicates. Data are provided as mean values +/- standard deviation. Source data are provided as a Source Data file.

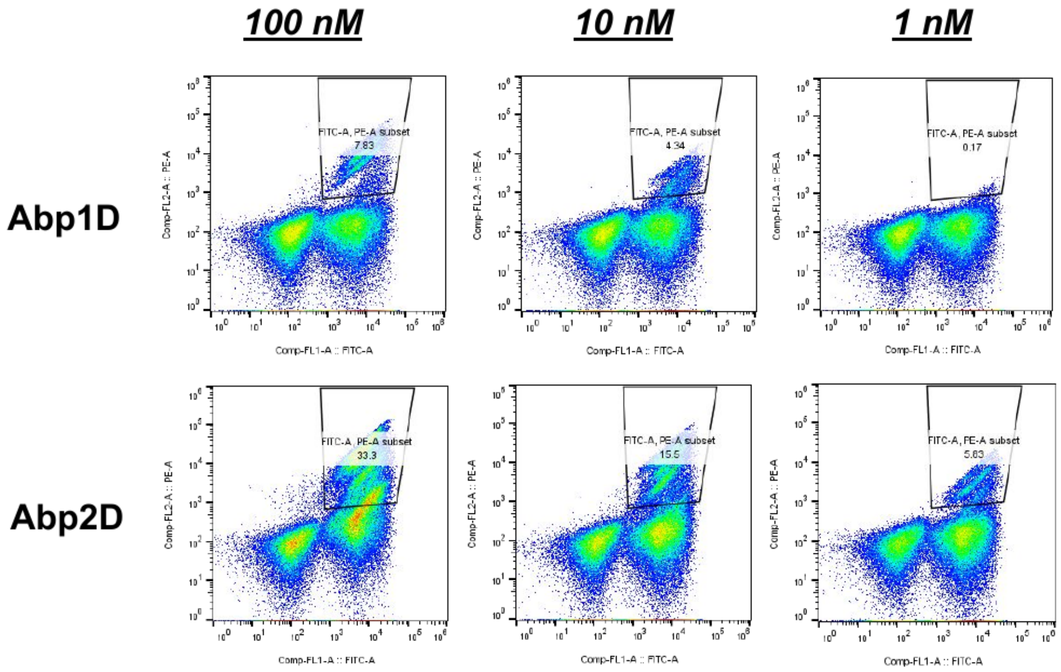


**Fig. S11. Titration sort results for yeast library and Abps.** FACS results for sort 3 of the yeast display experiment using 10-fold titrations of Abp1D and Abp2D. Data analyzed using FlowJo. Source data from FCS files are provided in Supplementary Data File 2.

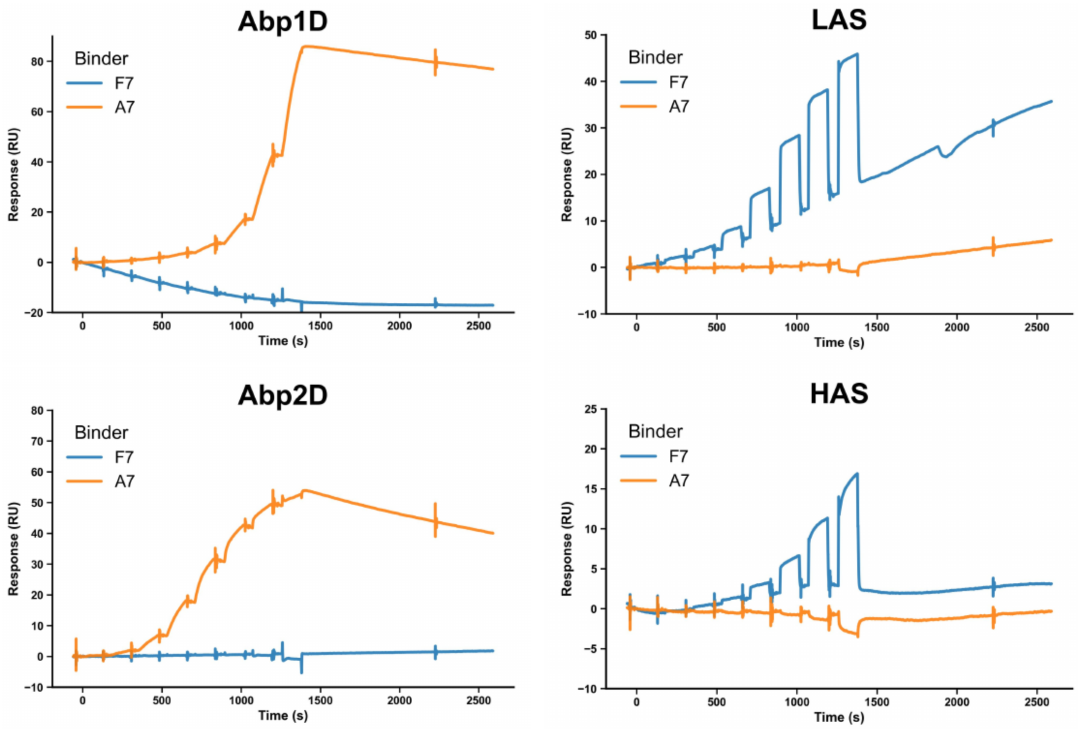


**Fig. S12.** ***In vitro* specificity test for adhesin minibinders.** SPR traces for Abp minibinder A7 and FimH minibinder F7 with Abp1D, Abp2D, FimH LAS, and FimH HAS. Source data are provided as a Source Data file.

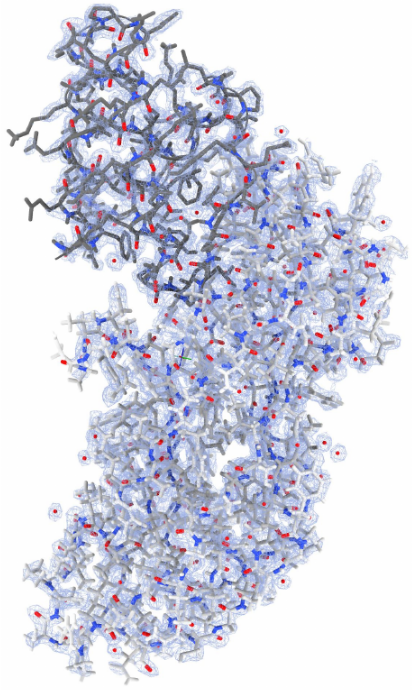

**Fig. S13. Electron density map of A7-Abp2D co-crystal.** 2mFo-DFc electron density maps contoured at 1 $\sigma$ shown in blue. A7 is shown in grey and Abp2D is shown in white.

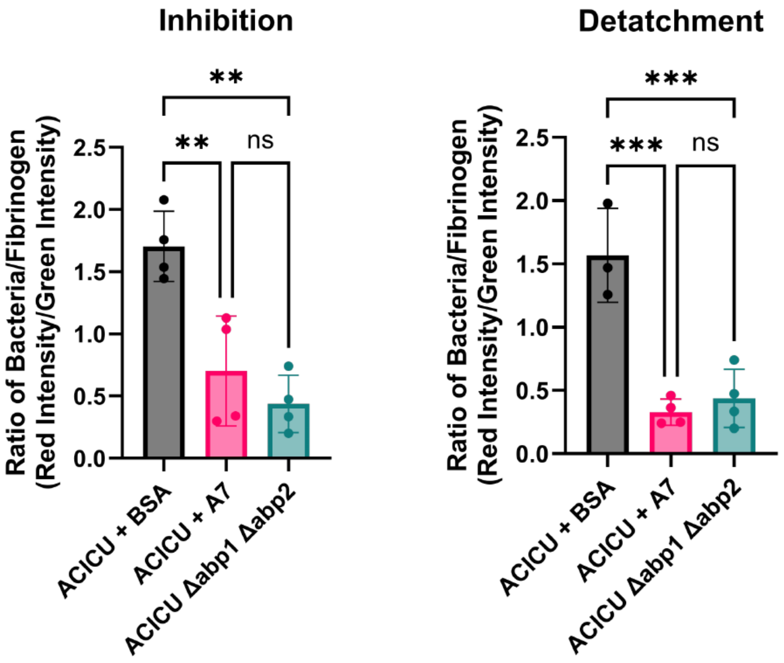


**Fig. S14.** Quantification of *A. baumannii* binding to treated catheters preincubated with 100 nM of A7 inhibitor (left) or applied to catheters after bacterial attachment (right). Normalized signal intensity of *A. baumannii* bacteria (red signal) over fibrinogen coating (green signal) per catheter. Inhibition: ACICU + BSA vs ACICU + A7 treatment p=0.0052. Inhibition: ACICU + BSA vs ACICU Δabp1 Δabp2 + BSA treatment groups p=0.0011. Detachment: ACICU + BSA vs ACICU + A7 treatment groups p=0.0004. Detachment: ACICU + BSA vs ACICU Δabp1 Δabp2 + BSA treatment groups p=0.0007. n=4 for all groups, except for ACICU + BSA detachment where n=3. Data are presented as mean values +/- standard deviation. One-way ANOVA with Tukey's multiple comparisons test. ***P $\leq$ 0.001, **P $\leq$ 0.01, *P $\leq$ 0.05. Source data are provided as a Source Data file.

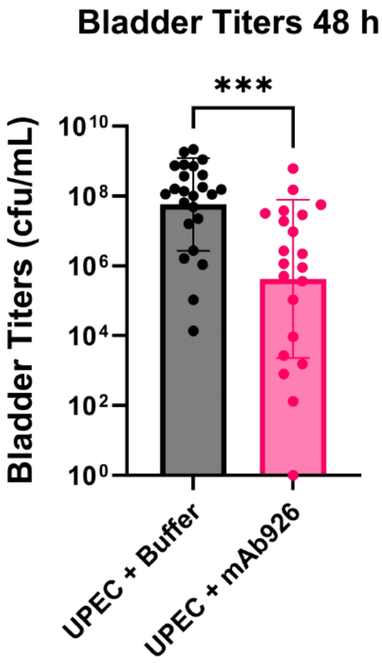


**Fig. S15.** Titers of bacteria 48 hours post-infection (hpi) in the bladder after treatment with buffer (grey, n=24), or monoclonal antibody (pink, n=21). UPEC + Buffer vs UPEC + F7 p=0.0001. Data are presented as mean values +/- SD of n biological replicates. Mann-Whitney U test with two-tailed P value. ***P $\leq$ 0.001, **P $\leq$ 0.01, *P $\leq$ 0.05. Each data point represents one mouse. n indicates the number of mice per treatment group. Source data are provided as a Source Data file.

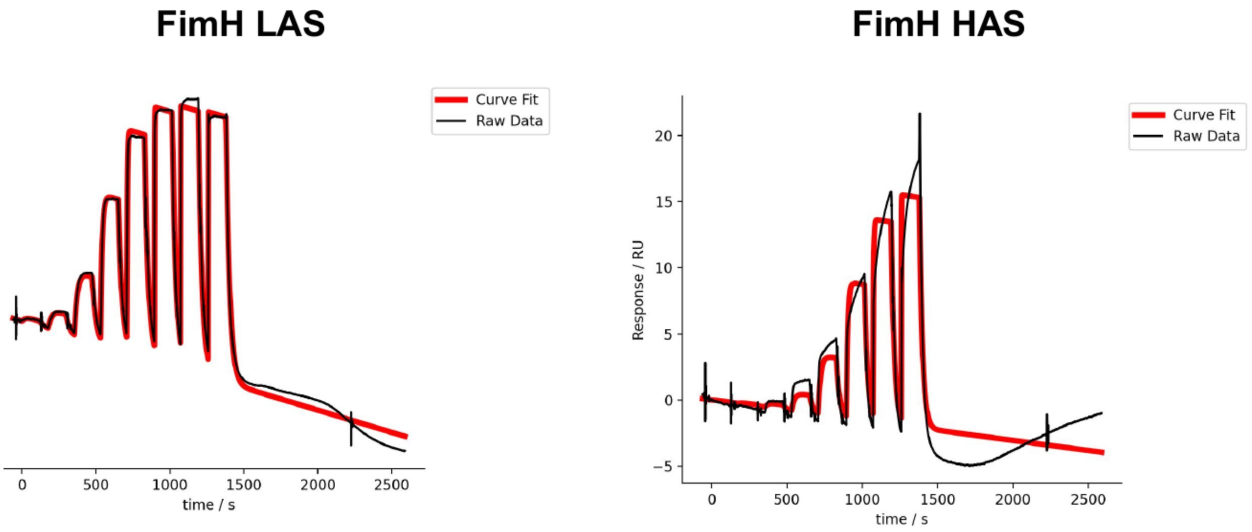


**Fig. S16. SPR traces of FimH minibinder C8.** Minibinder C8 binds with a higher affinity than F7 to both FimH LAS ($K_d$=15.0 nM) and FimH HAS ($K_d$=243 nM). Source data are provided as a Source Data file.

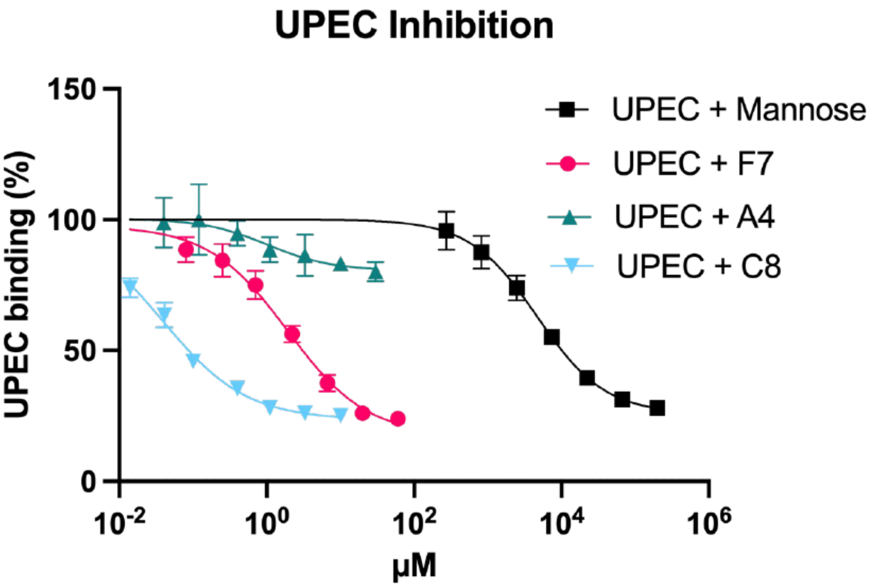


**Fig. S17. Inhibition ELISA of minibinder C8.** Inhibition ELISA results for minibinder F7 (pink; IC50=1.9 µM; 95% CI: 1.4-2.9 µM), noninhibitory minibinder A4 (teal), improved minibinder C8 (cyan; IC50-37 nM; 95% CI: 31-47 nM) and mannose (black). Each experiment includes three replicates. Data are presented as mean values +/- SD. Source data are provided as a Source Data file.

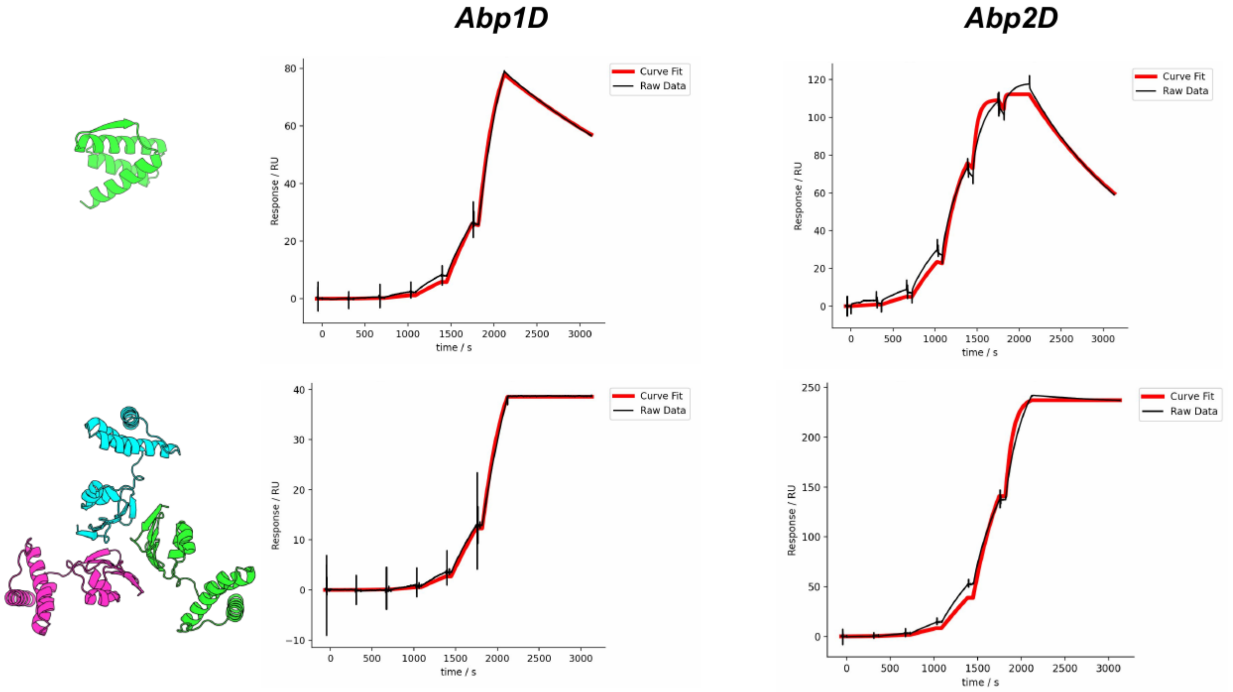


**Fig. S18. SPR traces of Abp Oligomer C11 and its parent: Abp minibinder A7.** Oligomer C11 binds with a higher affinity than its parent to both Abp1D ($K_d$=11 pM) and Abp2D ($K_d$=195 pM). Source data are provided as a Source Data file.

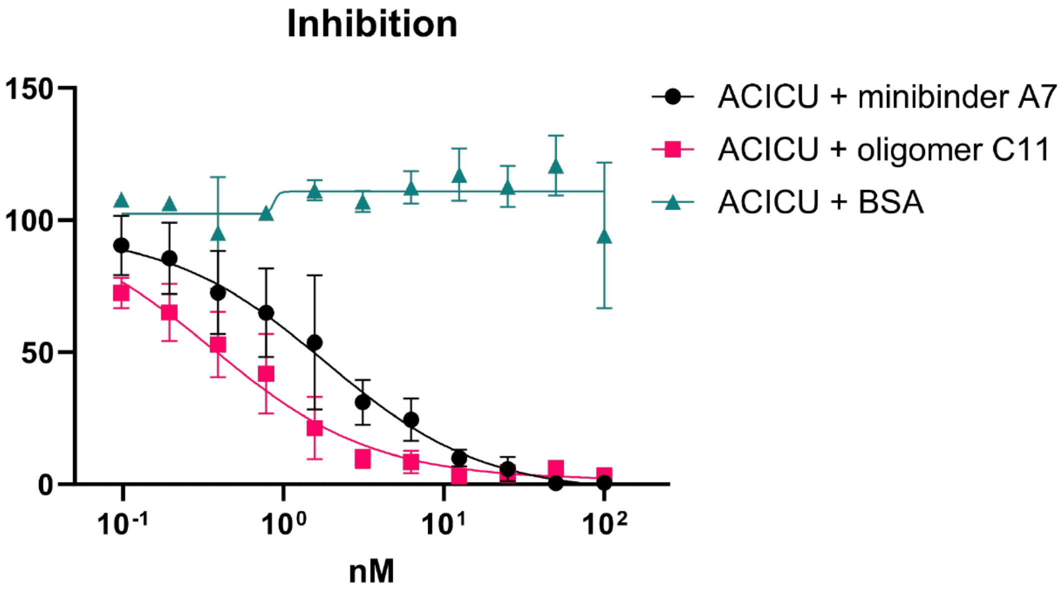


**Fig. S19. Inhibition ELISA results for Abp oligomer C11 and its parent: Abp minibinder A7.** ELISA results indicate that oligomerization improves the IC50 of inhibition of fibrinogen binding *in cellulo*. The data provided contains 3 technical replicates. Data are presented as mean values +/- SD. Source data are provided as a Source Data file.

### Supplementary Tables

**Table S1. $K_d$ estimates of FimH minibinders during initial screen.** This table includes the kinetic profiles ($K_d$s to LAS and HAS) of FimH minibinders that displayed affinity for either the HAS or LAS.

| Binder_ID | LAS – $K_d$ (nM) | HAS – $K_d$ (nM) |
|---|---|---|
| A3 | 5000 | 3700 |
| A4 | 2000 | 2500 |
| B1 | 266 | N.D. |
| B2 | 634 | N.D. |
| C10 | 300 | N.D. |
| C3 | 437 | N.D. |
| C5 | 2000 | 2200 |
| C7 | 2000 | N.D. |
| D9 | 3500 | 4700 |
| E10 | 4300 | 4200 |
| E8 | N.D. | 4800 |
| F1 | 939 | 729 |
| F4 | 149 | N.D. |
| **F7** | **100** | **N.D.** |
| G12 | 132 | N.D. |
| H9 | 198 | N.D. |

N.D.: Binding was not detected. The $K_d$ estimate is larger than the highest screened concentration (5 µM).
Bold: the design in bold was selected for further characterization.

**Table S2. Designed protein sequences.**

| Target | Binder | Binder sequence |
|---|---|---|
| FimH | FimH minibinder F7 | MAEKEAALTAAADGTVAALAAGNIGVDYARYRRKALVAYAKKEGLPQAVIDAVTARLDAAIAAAEAA |
| FimH | FimH minibinder C8 | MEEKIKAAKEAADGTVAALAAGNIGVDYARYYKKALVAWMKKQGLPQEVIDEVTAKLDAAIAAAEAA |
| Abp1D, Abp2D | Abp minibinder A7 | EKSYEEAVLEANKLIESGAPDEEVEKATKYALDKYAASIGLSVVEYPPLETLKEFVTKEAAKIRAA |
| Abp1D, Abp2D | Abp oligomer C11 | MKVYEFPYPETGKKIIVIQGEKNIVIVVGNTAVVYYEGKWTYKENVTEEDIEKAKTEEGAKELAKGSGEKSYEEAVLEANKLIESGAPDEEVEKATKYALDKYAASIGLSVVEYPPLETLKEFVTKEAAKIRAA |

**Table S3. Data collection and refinement statistics for crystal structures.** Statistics for the highest-resolution shell are shown in parentheses.

| | F7-FimH (PDB Code: 9Q1V) | A7-Abp2D (PDB Code: 9Q1H) |
|---|---|---|
| Resolution range | 42.22 – 1.75 (1.81 – 1.75) | 49.60 – 1.35 (1.398 – 1.35) |
| Space group | P $2_1$ | P $2_1$ $2_1$ $2_1$ |
| Unit cell | 50.97, 52.39, 72.87; 90, 101.80, 90 | 25.81, 52.46, 152.12; 90, 90, 90 |
| Unique reflections | 48567 (5071) | 46366 (4424) |
| Multiplicity | 6.9 (7.2) | 2.0 (2.0) |
| Completeness (%) | 90.00 (95.91) | 99.43 (97.34) |
| Mean I/sigma(I) | 3.99 (1.30) | 17.87 (2.02) |
| Wilson B-factor | 15.58 | 13.46 |
| R-merge | 0.2215 (1.516) | 0.02098 (0.3197) |
| R-pim | 0.0900 (0.603) | 0.02098 (0.3197) |
| $CC_{1/2}$ | 0.995 (0.402) | 0.999 (0.759) |
| Reflections used in refinement | 34326 (3632) | 46363 (4424) |
| R-work | 0.2330 (0.3709) | 0.1648 (0.1912) |
| R-free | 0.2630 (0.3957) | 0.1971 (0.2624) |
| Number of non-hydrogen atoms | 3689 | 1834 |
| macromolecules | 3453 | 1754 |
| solvent | 236 | 80 |
| Protein residues | 465 | 229 |
| RMS(bonds) | 0.004 | 0.009 |
| RMS(angles) | 0.66 | 0.96 |
| Ramachandran favored (%) | 97.81 | 98.67 |
| Ramachandran allowed (%) | 2.19 | 1.33 |
| Ramachandran outliers (%) | 0.00 | 0.00 |
| Average B-factor | 22 | 18 |
| macromolecules | 22 | 16 |
| solvent | 25 | 22 |

**Table S4. $K_d$ estimates of Abp minibinders during initial screen.** This table includes the kinetic profiles ($K_d$s to Abp1D and Abp2D) of Abp minibinders that were enriched during yeast surface display and expressed sufficiently for downstream characterization as measured by SPR. Minibinders that exhibited undetectable binding to both Abp1D and Abp2D were excluded for clarity.

| Binder_ID | Abp1D – $K_d$ (nM) | Abp2D – $K_d$ (nM) |
|---|---|---|
| A4 | 40 | 559 |
| A6 | 3900 | P.F. |
| **A7** | **67** | **4** |
| A8 | 2030 | P.F. |
| A12 | N.D. | 235 |
| B1 | P.F. | 48 |
| B4 | N.D. | 786 |
| B5 | 987 | 1010 |
| B6 | 1260 | 336 |
| **B7** | **101** | **16** |
| B8 | 3830 | 201 |
| B9 | 669 | 187 |
| B12 | N.D. | 657 |
| C1 | 1240 | 624 |
| C6 | N.D. | 950 |
| **C7** | **6** | **124** |

N.D.: Binding was not detected. The $K_d$ estimate is larger than the highest screened concentration (5 µM). P.F.: the binding data could not be fit well to accurately estimate a $K_d$. Bold: the designs in bold exhibited $K_d$s to both Abp1D and Abp2D at or below 1 µM.
